# Cortical language areas are coupled via a soft hierarchy of model-based linguistic features

**DOI:** 10.1101/2025.06.02.657491

**Authors:** Ahmad Samara, Zaid Zada, Tamara Vanderwal, Uri Hasson, Samuel A. Nastase

**Affiliations:** University of British Columbia, Vancouver, BC, Canada; Princeton Neuroscience Institute, Princeton University, Princeton, NJ, USA; Department of Psychology, Princeton University, Princeton, NJ, USA; BC Children’s Hospital Research Institute, Vancouver, BC, Canada; University of Southern California, Los Angeles, CA, USA

**Keywords:** encoding models, functional connectivity, language network, large language models (LLMs), narrative comprehension, naturalistic paradigm

## Abstract

Natural language comprehension is a complex task that relies on coordinated activity across a network of cortical regions. In this study, we propose that regions of the language network are coupled to one another through subspaces of shared linguistic features. To test this idea, we developed a model-based connectivity framework to quantify stimulus-driven, feature-specific functional connectivity *between* language areas during natural language comprehension. Using fMRI data acquired while subjects listened to spoken narratives, we tested three types of features extracted from a unified neural network model for speech and language: low-level acoustic embeddings, mid-level speech embeddings, and high-level language embeddings. Our modeling framework enabled us to quantify the proportions of stimulus features driving connectivity between regions: early auditory areas were coupled to intermediate language areas via lower-level acoustic and speech features; in contrast, higher-order language and default-mode regions were predominantly coupled through more abstract language features. We observed a clear progression of feature-specific connectivity from early auditory to lateral temporal areas, advancing from acoustic connectivity to speech- and finally to language-driven connectivity. Our findings suggest that higher-order language areas are coupled along increasingly higher level, more contextualized language features.

## Introduction

Language is a fundamental part of everyday human behavior that allows us to communicate complex ideas from one person to another. Our ability to understand language is remarkably efficient given the complexity of the task. As we listen to spoken language, we rapidly convert acoustic signals into words, link words into complex grammatical structures, and integrate all of these patterns into a holistic understanding of the discourse or situation (Christiansen & Chater, 2016). In most cases, we do this effortlessly. Language comprehension emerges from the coordinated activity of a number of different brain areas. A large body of research on the neurobiology of language has identified a highly interconnected network of cortical regions that are selectively engaged during language processing (Hickok & Poeppel, 2007; Friederici, 2011; Price, 2012; Fedorenko et al., 2024). How these language regions coordinate with one another to support efficient language comprehension, however, remains incompletely understood.

Regions of the language network are both structurally interconnected (Catani et al., 2005; Duffau, 2008; Saur et al., 2008; Friederici, 2009; Turken & Dronkers, 2010; Dick et al., 2014) and functionally integrated (Hampson et al., 2002; Lohmann et al., 2010; Lee et al., 2012; Tomasi & Volkow, 2012; Blank et al., 2014; Tie et al., 2014; Zhu et al., 2014; McAvoy et al., 2016; Kong et al., 2021; Du et al., 2024; Salvo et al., 2025). Recent work, for example, has shown that language areas are functionally interconnected even during rest and non-linguistic tasks (Braga et al., 2020; Shain & Fedorenko, 2025). However, traditional within-subject functional connectivity (WSFC) metrics cannot distinguish between intrinsic and extrinsic (i.e., stimulus-driven) co-fluctuations between regions. A between-subject approach is becoming increasingly common in naturalistic neuroscience, whereby intersubject correlation (ISC) analyses isolate stimulus-evoked activity within a brain region (Hasson et al., 2004; Nastase et al., 2019). Following the logic of ISC, intersubject functional connectivity (ISFC) has been used to isolate stimulus-driven connectivity between regions in response to naturalistic stimuli such as spoken narratives (Simony et al., 2016). However, ISFC captures the stimulus-driven components of functional connectivity in a data-driven fashion that is agnostic to the content of the stimulus shared between regions: any features of the stimulus can drive connectivity between any two regions. ISFC can tell us *where* and *how much* connectivity is driven by the stimulus, but not *which* stimulus features are driving the connectivity.

How can we begin to unravel *what* linguistic features are shared across different language regions? A growing body of recent work indicates that different kinds of linguistic structures (e.g., syntax and semantics) appear to be co-localized across language areas (Fedorenko et al., 2012, 2016; 2020; Wehbe et al., 2014; Blank et al., 2016; Nelson et al., 2017; Caucheteux et al., 2021; Reddy & Wehbe, 2021; Toneva et al., 2022a; Kumar, Sumers et al., 2024; Shain et al., 2024), and most language areas display similar response profiles to a variety of language tasks (Fedorenko et al., 2024). These observations create a certain tension: surely, different language areas contribute differently to the overall circuit, but why do we observe such overwhelming functional similarity across regions? From a computational perspective, one possible explanation for this tension is that different areas of the language network may interact through a “communication subspace” of shared features (Semedo et al., 2019). For example, in visual areas, one region is coupled to another via fluctuations along a subset of dimensions in the population-level activity space (Semedo et al., 2019; Kohn et al., 2020; MacDowell et al., 2025). Perhaps different regions of the language network rely on a shared, multidimensional embedding space of linguistic features to efficiently coordinate their contributions to the network as a whole. Interestingly, modern large language models (LLMs) appear to rely on a similar geometric mechanism of connectivity: the circuits across layers of an LLM interact with one another by gradually refining linguistic representations as they proceed through a high-dimensional embedding space shared across layers (Elhage et al., 2021). This so-called “residual stream” serves as a communication channel across layers and is thought to be critical to the capacity of LLMs to capture the rich, content-specific structure of natural language.

In this study, we hypothesized that different regions of the language network are coupled to one another via a multidimensional space of linguistic features. This hypothesis is inspired by modern neural network models for speech and language, which integrate phonemic, syntactic, and semantic features into a unified neural population code (e.g., Radford et al., 2023), and construct increasingly refined representations by modifying a high-dimensional embedding space that is shared from layer to layer (Elhage et al., 2021). In a similar way, we hypothesize that two regions of the language network may harmonize their contributions to language comprehension via a shared subspace of linguistic features. Our hypothesis yields two predictions. First, functional connectivity between one language region and another should result from moment-to-moment, stimulus-driven covariation along a shared subset of linguistic features. Second, as we proceed along the cortical processing hierarchy, connectivity should be driven by increasingly abstract, more contextualized linguistic features. Given the mixed selectivity (or “polysemanticity”) of neural population codes (Fusi et al., 2016; Elhage et al., 2022; Bricken et al., 2023), we expect some low- and mid-level features to be retained even as other features become increasingly complex; we refer to this as a “soft hierarchy”.

To test these predictions, we developed a novel model-based framework for quantifying stimulus-driven, feature-specific co-fluctuations in neural activity between one region and another. This model-driven framework provides a theoretical advance over content-agnostic metrics of functional connectivity (both WSFC and ISFC) by allowing us to test explicit, feature-specific models of the functional connectivity between language areas. To this end, we decomposed two spoken stories into low-level acoustic, mid-level speech, and high-level language features based on embeddings extracted from the Whisper speech and language model (Radford et al., 2023). We tested these model features against naturalistic fMRI data acquired while subjects listened to the same spoken stories. Our findings reveal a soft processing hierarchy where language areas are coupled along shared acoustic, speech, and language features, and connectivity from lower- to higher-order areas is driven by progressively more contextualized linguistic features.

## Results

To investigate how regions of the language network coordinate their activity during naturalistic language comprehension, we developed a model-based framework that quantifies stimulus-driven, feature-specific functional connectivity between brain areas. Before reporting our core results, we first develop the theoretical motivation for our approach and validate our models within each brain region. ISC analysis (Hasson et al., 2004; Nastase et al., 2019) has been used, initially, to isolate the stimulus-driven component of neural activity within a given brain region (**Fig. 1a**). In separate subjects (scanned at separate times), the stimulus is the only shared factor that could drive shared activity. However, ISC analysis is data-driven and content-agnostic: it can tell us *where* and *how much* activity is driven by the stimulus, but it cannot determine *what* stimulus features drive neural activity (Zada et al., 2024). To quantify *what* stimulus features are driving activity in a given brain region, we use parcel-wise encoding models to quantify which explicit linguistic features are encoded in brain activity (Wu et al., 2006; Nalesaris et al., 2011; Huth et al., 2016; de Heer et al., 2017; Kell et al., 2018; Dupré la Tour, Visconti di Oleggio Castello et al., 2025; **Fig. 1b**).

**Fig. 1.**
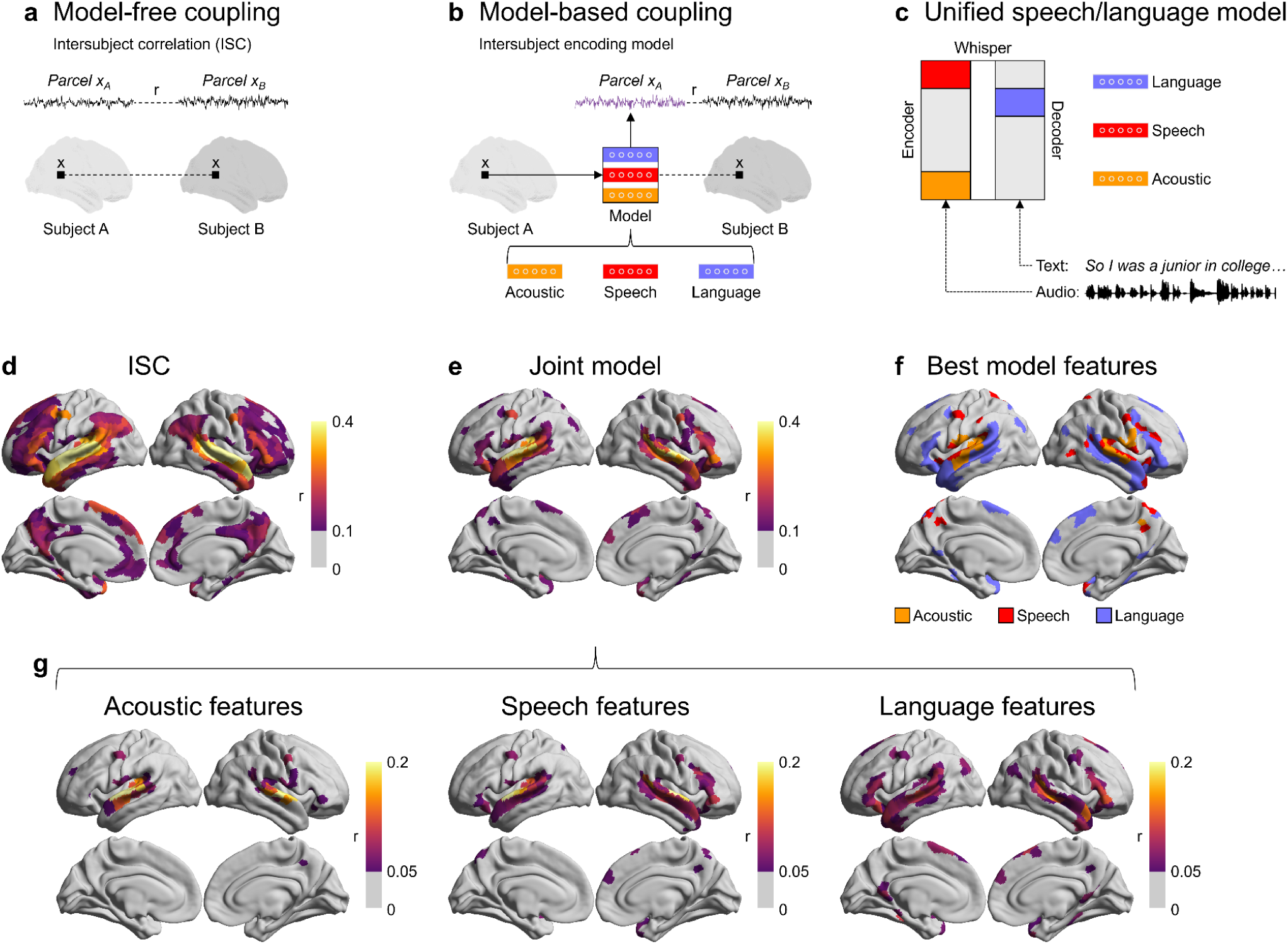
Modeling stimulus-driven, feature-specific neural activity during natural language comprehension. **(a)** Intersubject correlation is a data-driven, model-free method for quantifying the stimulus-driven component of neural activity *within* a given region. **(b)** To quantify the activity driven by specific stimulus features, we construct parcel-wise encoding models using explicit linguistic features extracted from a computational model. Encoding models are evaluated within a given brain region by correlating model-predicted activity with the actual activity in a left-out subject or group of subjects (in the same way as ISC analysis). **(c)** We extract three types of linguistic features from a unified transformer-based speech and language model called Whisper: acoustic (orange), speech (red), and language (blue) feature representations (i.e., embeddings) of the stimulus. The schematic shows a more detailed version of the model depicted in **(b)**. For more details on the encoding model approach, refer to **Fig. S1**. Dashed lines indicate correlation; solid lines indicate model input/output. **(d)** We computed ISC within 1,000 cortical parcels for two story-listening fMRI datasets. Parcels with significant ISC (p < .05) were further thresholded at 0.10 for visualization. Encoding models were estimated across all three feature bands using banded ridge regression. **(e)** Performance for the joint model (i.e., encoding model fit jointly using all three feature bands) and thresholded in the same way **(e)**. **(f)** The best-performing feature band was identified for each parcel. **(g)** The joint model performance can be decomposed into feature-specific performance values.

To quantify the linguistic features of the stimulus, we utilized a state-of-the-art transformer-based neural network model for speech recognition, known as Whisper (Radford et al., 2023). We elected to use Whisper because it is a unified acoustic-to-speech-to-language model that learns how to map the acoustic signals of natural speech into language representations useful for natural language tasks like next-word prediction and transcription (**Fig. 1c**). The “encoder” component of the model learns to extract linguistic features from acoustic inputs, whereas the “decoder” component extracts linguistic features from text inputs. We extracted three types of linguistic features from Whisper: (1) low-level “acoustic” embeddings from the input to the transformer stack of the encoder: (2) mid-level “speech” embeddings from the final layer of the encoder; and (3) high-level, more abstract “language” embeddings from a late-intermediate layer of the decoder. While the acoustic embeddings are non-contextual and pool over very brief windows (the duration of a word), the speech embeddings are contextualized by preceding sounds, and the language embeddings are further contextualized by preceding words. These three sets of embeddings capture increasingly abstract and contextual features of spoken language that the model uses to perform natural language tasks (Goldstein et al., 2025). All three embeddings were of the same dimensionality (1,024 dimensions). To ensure that our results are not specific to Whisper, we also performed the main analyses using alternative acoustic, speech, and linguistic features extracted from two other models: acoustic and speech embeddings from the HuBERT speech model (Hsu et al., 2021) and linguistic features from the Gemma language model (Riviere et al., 2024).

We evaluated our models against fMRI data comprising *N* = 46 participants, each of whom listened to two different ∼13-minute spoken stories (Nastase et al., 2021). To reduce computational demands, we first reduced the voxel-level time series into 1,000 parcel-level time series derived from a functional atlas (Schaefer et al., 2018). We used banded ridge regression to fit parcel-wise encoding models jointly across all three sets of features (acoustic, speech, and language embeddings; Dupré la Tour et al., 2022). This allows all three types of linguistic features to compete for variance in the neural activity fairly. To test the alignment of linguistic features with human brain activity, we generated predictions from each of the three feature bands for the left-out data and computed the correlation between the model-predicted and actual brain activity at each parcel. We also computed the joint model performance, which corresponds to the sum of the predictions for each of the three feature bands. To more closely match the formulation of ISC, we estimated encoding models within each subject and evaluated their performance across subjects by computing the correlation of model-based predictions from one subject with the average actual time series across all other subjects. While this intersubject encoding approach differs from much prior work (e.g., Huth et al., 2016; Schrimpf et al., 2021; Goldstein et al., 2022, 2025; cf. Van Uden, Nastase et al., 2018; Nastase et al., 2020b; Zada et al., 2024), the focus of our analyses is ultimately on what features are shared between regions (not between subjects). To further ensure the generalizability of our findings, we estimated all models within one story and evaluated their performance in predicting the other story (and vice versa; **Fig. S1**). Overall, this modeling framework allows us to quantify how well the model features linearly align with human neural activity during complex, naturalistic language comprehension.

### Modeling the soft hierarchy of linguistic features across the language network

We first localized language areas involved in processing the story stimuli using a conventional ISC analysis, which identifies parcels where neural activity is synchronized to the stimulus (**Fig. 1a**). ISC analysis revealed a large-scale cortical network for spoken narrative comprehension comprising low-level auditory areas, language areas, and higher-level default-mode areas (**Fig. 1d**) (Nastase et al., 2021). Next, we evaluated the joint encoding model performance combined across all three feature bands from the Whisper model using our intersubject encoding approach (**Fig. 1b**). The resulting joint model predicted neural activity across much of the language network, including parcels in frontotemporal language regions, as well as in posterior medial cortex (PMC), superior frontal language area (SFL), dorsomedial prefrontal cortex (dmPFC; **Fig. 1e,g**). The joint model performance map captures a subset of regions identified by the ISC analysis, suggesting that ISC identifies certain stimulus features of spoken stories that are not represented in the embeddings extracted from the Whisper model.

We then evaluated predictions derived separately from each of the three feature bands. By visualizing the top-performing feature at every parcel (**Fig. 1f**), we identified a coarse processing hierarchy where acoustic features dominate in superior temporal auditory areas, speech features are more sparsely distributed along temporal cortex and higher-level areas, and the language features dominate in lateral temporal areas (both anterior and posterior) and inferior frontal gyrus (IFG). The acoustic model performance was largely confined to the early auditory cortex (EAC) and superior temporal gyrus (STG), with punctate clusters in middle frontal gyrus (MFG; **Fig. 1g**). Speech model performance extended from EAC and STG anterolaterally to superior temporal sulcus (STS), and included portions of right IFG (**Fig. 1g**). Language embeddings predicted a broader array of language regions, including parcels in both anterior and posterior lateral temporal cortex, IFG, as well as PMC, SFL, dmPFC (**Fig. 1g**). These results are generally consistent with prior work using encoding models to map linguistic features onto cortical activity (e.g., Wehbe et al., 2014; Huth et al., 2016; de Heer et al., 2017; Kell et al., 2018; Goldstein et al., 2025). In line with the notion of a soft hierarchy, we found significant overlap among the feature-specific model performance maps, suggesting that many cortical areas encode mixed acoustic, speech, and language features (**Fig. 1g**).

### Modeling stimulus-driven, feature-specific connectivity between language areas

Next we turned to the core question of this manuscript: How can we quantify the stimulus-derived linguistic features driving connectivity *between regions* of the language network? First, by extending the same logic as ISC, we can use ISFC analysis to quantify the stimulus-driven *connectivity* between brain areas, effectively filtering out idiosyncratic noise and the intrinsic fluctuations that play a large role in traditional WSFC (Simony et al., 2016; **Fig. 2a**). ISFC analysis yields a parcel-by-parcel matrix of stimulus-driven connectivity values between pairs of brain regions. The diagonal of this matrix corresponds to the within-region ISC values (**Fig. 1a**). Similar to both ISC analysis and WSFC analysis, ISFC analysis is a data-driven method that indicates *where* and *how much* stimulus-driven connectivity exists between two regions; however, it does not reveal *what* features of the stimulus drive that connectivity.

**Fig. 2.**
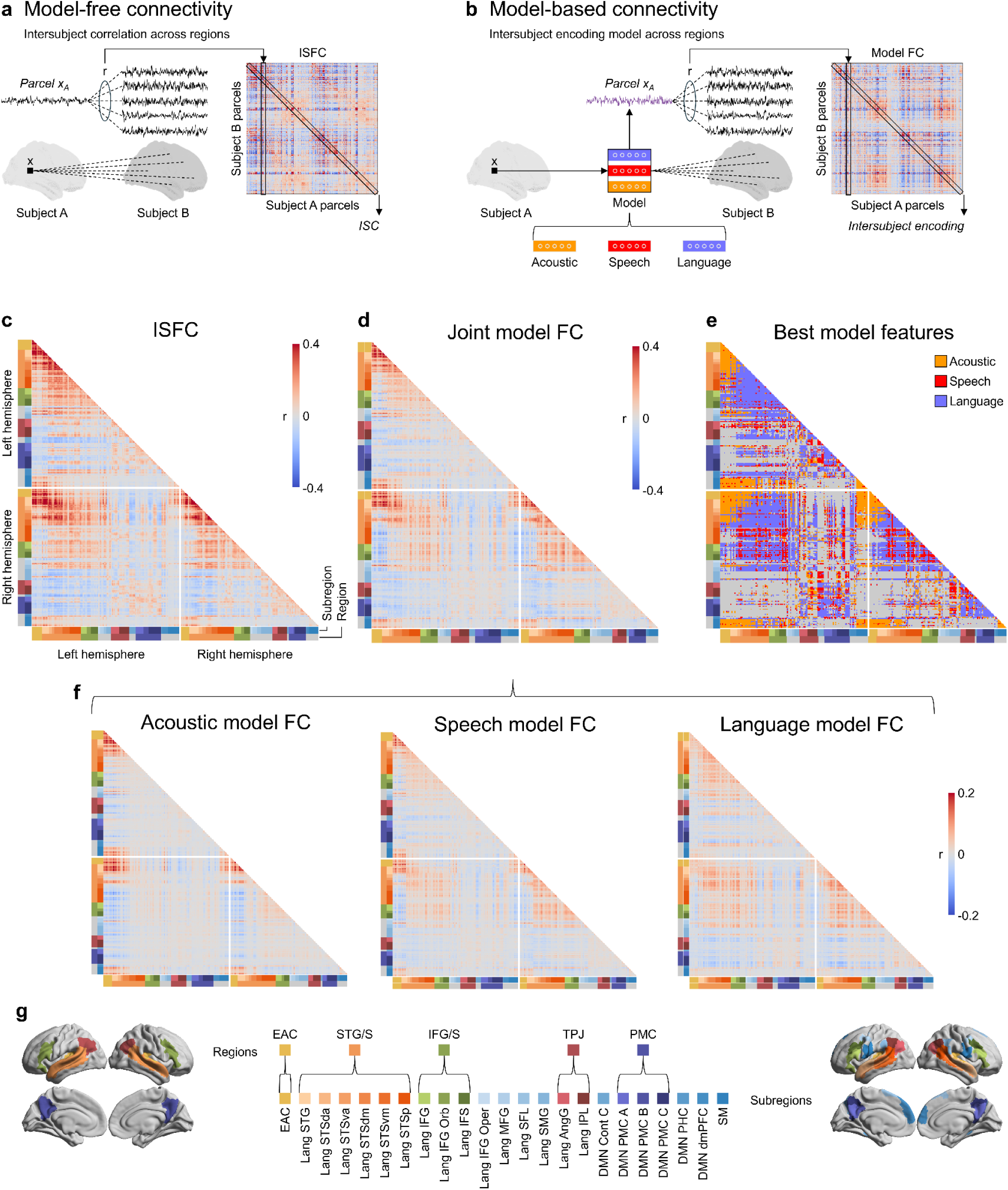
Modeling stimulus-driven, feature-specific functional connectivity during natural language comprehension. **(a)** Following the logic of ISC, intersubject functional connectivity (ISFC) quantifies the stimulus-driven connectivity between brain areas in a data-driven, model-free fashion. The diagonal of the ISFC matrix corresponds to the within-parcel ISC values. **(b)** To quantify feature-specific model-based connectivity, we evaluate encoding models based on the three sets of linguistic features extracted from Whisper *across* pairs of parcels. The diagonal of the resulting model-based connectivity matrix corresponds to the within-parcel intersubject encoding performance values. Dashed lines indicate correlation; solid lines going into and out of the model indicate model input/output. **(c)** A parcel-by-parcel ISFC matrix was computed for all pairs of parcels within language areas. The joint model-based connectivity matrix **(d)**, as well as feature-specific model connectivity (model FC) matrices **(f)** were computed in the same way as the ISFC matrix for each subject and were averaged across subjects for visualization. Feature-specific model FC matrices were computed based on acoustic, speech, and language embeddings extracted from the Whisper model. **(e)** The best-performing feature space at each edge was chosen as the one with the correlation value closest to the original ISFC value at that edge. The best feature map is thresholded to include only edges that are positive in the original ISFC matrix. **(g)** All connectivity matrices are visualized for a set of 280 parcels spanning functionally defined language regions (see *Regions of Interest* section under *Methods* for details on how these regions were defined).

To quantify *what* stimulus features are shared between different brain regions, we again used linguistic embeddings extracted from the Whisper model. We use the same parcel-wise encoding models trained within each parcel from the previous section. In our model-based connectivity analysis, we now evaluate these models in terms of how well their predictions generalize to *other parcels* in the same fashion as the ISFC analysis. We generate model-based predictions for one parcel, then correlate these predicted time series with the average actual time series of the remaining subjects across all pairs of parcels (**Fig. 2b**). This model-based functional connectivity analysis results in a parcel-by-parcel matrix of feature-specific connectivity values between pairs of parcels (Toneva et al., 2022b; Meschke et al., 2023; Zada et al., 2024). In this case, the diagonal of the connectivity matrix corresponds to the within-parcel intersubject encoding model performance (**Fig. 1b**). The use of encoding models effectively filters the connectivity between regions based on what can be captured by an explicit feature space. This analysis quantifies the extent to which activity in different regions of the language network is driven by the same set of linguistic features occurring at the same time in the stimulus.

We first visualized the conventional ISFC matrix (**Fig. 2c**) and the joint model-based connectivity matrix based on the combination of all three sets of linguistic features (**Fig. 2d**). The joint model connectivity matrix appears to recapitulate some but not all of the stimulus-driven connectivity structure in the ISFC matrix, albeit with lower correlation values. We also constructed model-based connectivity matrices based on predictions derived from each of the three types of linguistic features. These parcel-by-parcel model-based connectivity matrices are complex, and much of the rest of our analyses aim to interpret these feature-specific networks. When examining the best-performing feature band at each connection, we see relatively focal connectivity for the acoustic embeddings, sparse connectivity for the speech embeddings, and more widespread connectivity for the language embeddings (**Fig. 2e**). The acoustic model best captured shared connectivity within EAC and between EAC and STG/S parcels. In contrast, the language model outperformed the other two models in capturing shared connectivity within and between STG/S and IFG/S parcels. Model performance was comparatively lower in the temporoparietal junction (TPJ) and PMC regions, with mixed results across models. Overlapping patterns of connectivity across matrices for the three types of stimulus features suggest that many edges (i.e, connections between parcels within or across regions) may be captured by overlapping sets of features (**Fig. 2f**). To simplify visualizing the connectivity matrices, we focused on a subset of 280 cortical parcels spanning 23 regions of interest (**Fig. 2g**) associated with language comprehension based on the ISC results (**Fig. 1d**). These model-based functional connectivity matrices serve as the basis for all of the subsequent analyses in the manuscript.

### Language regions are coupled via distinct and overlapping subspaces of linguistic features

We now focus on the cortical hierarchy for language comprehension and isolate components of network connectivity driven by specific stimulus features. We quantified the feature-specific model-based connectivity values within and between eight regions of interest ranging from low- to high-level areas for spoken narrative comprehension: EAC, STG/S, IFG/S, TPJ angular gyrus (TPJ AngG), TPJ inferior parietal lobule (TPJ IPL), PCM A, PMC B, and PMC C (**Fig. 3a**). We first examined the average local connectivity across all edges connecting parcels *within* a given region (**Fig. 3b**). We found that model connectivity among parcels within EAC was best predicted by acoustic (A) features, followed by speech (S) features, then language (L) features (A vs. S: t = 9.04, p_FDR_ < .001; S vs. L: t = 11.76, p_FDR_ < .001; A vs. L: t = 16.01, p_FDR_ < .001). The opposite was true for connectivity among parcels within STG (A vs. S: t = -8.21, p_FDR_ < .001; S vs. L: t = -7.21, p_FDR_ < .001; A vs. L: t = -11.16, p_FDR_ < .001), IFG (A vs. S: t = -11.09, p_FDR_ < .001; S vs. L: t = -4.25, p_FDR_ < .001; A vs. L: t = -11.56, p_FDR_ < .001), and appears to be the case in several other higher-level areas for language and narrative comprehension: the language embeddings captured the largest proportion of connectivity, followed by the speech embeddings, then the acoustic embeddings. This suggests that nearby parcels within a given language region tend to co-fluctuate along a shared set of linguistic features. Early auditory areas are most strongly coupled along acoustic and speech features. In contrast, parcels within higher-level areas, including language areas and default-mode areas, are coupled according to higher-level language features.

**Fig. 3.**
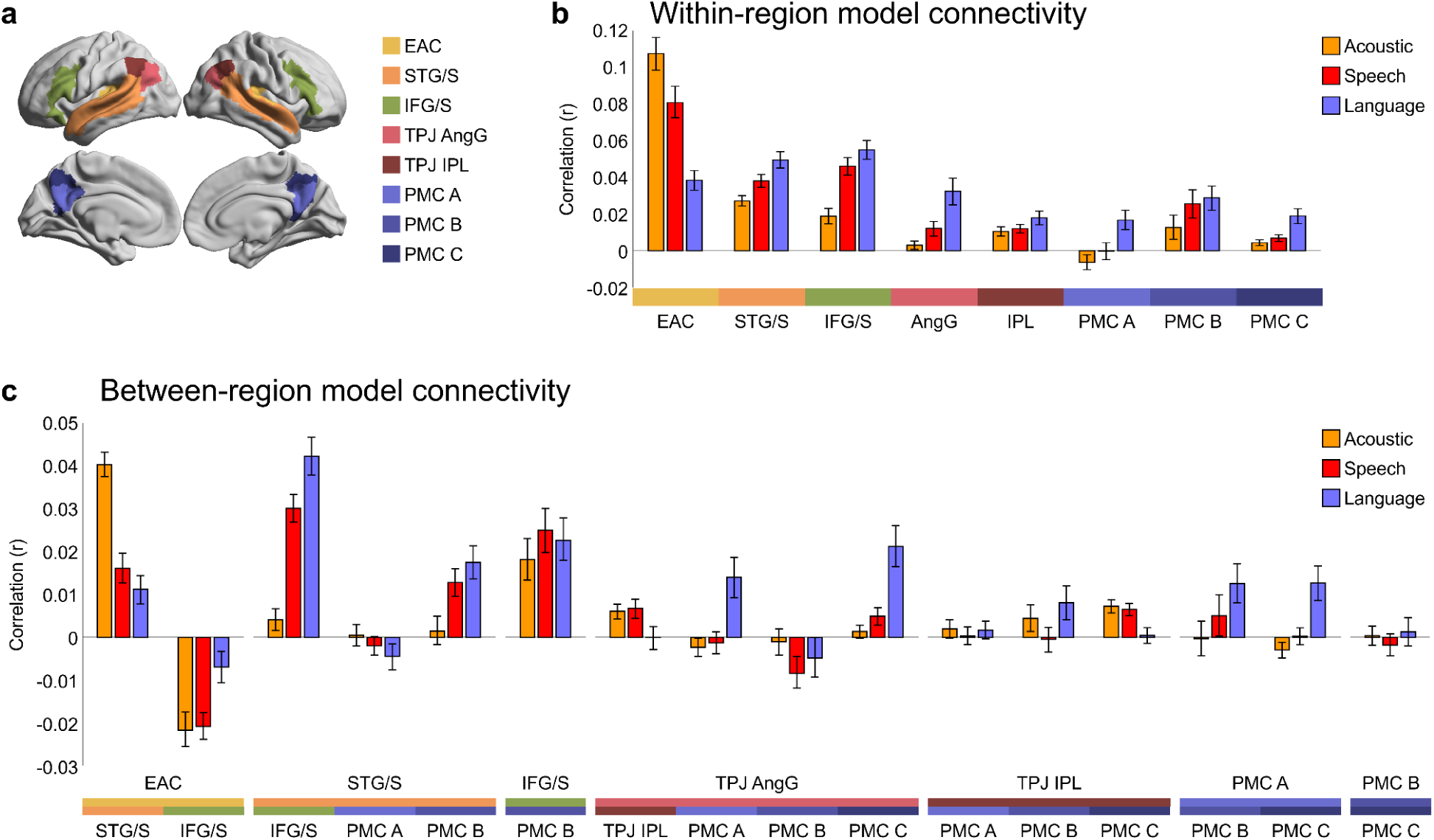
Feature-specific model-based connectivity within and between language regions. **(a)** We focused on eight language-related regions (comprising 207 parcels) that previously showed strong ISC during story listening: early auditory cortex (EAC), superior temporal gyrus and sulcus (STG/S), inferior frontal gyrus and sulcus (IFG/S), temporoparietal junction angular gyrus and inferior parietal lobule (TPJ AngG and TPJ IPL), and posterior medial cortex A, B, and C (PMC A, PMC B, and PMC C). **(b)** Feature-specific model connectivity values were averaged across parcel pairs within each language region. **(c)** Feature-specific model connectivity values were averaged across parcel pairs connecting different language regions. Between-region model connectivity results are shown only for region pairs with positive ISFC values. Error bars indicate bootstrap 95% confidence intervals. See **Fig. S4** for the same results plotted against the ISFC and joint model performance values.

We next examined feature-specific model-based connectivity *across* different regions of the language network (**Fig. 3c**). We found that the acoustic embedding best captured the connectivity between EAC and STG/S regions (A vs. S: t = 12.31, p_FDR_ < .001; S vs. L: t = 2.77, p_FDR_ = 0.009; A vs. L: t = 12.21, p_FDR_ < .001). The edges connecting STG/S and IFG, on the other hand, were dominated by the language embeddings, with speech embeddings capturing a large but secondary portion of variance (A vs. S: t = -18.50, p_FDR_ < .001; S vs. L: t = -6.79, p_FDR_ < .001; A vs. L: t = -17.48, p_FDR_ < .001). We observed a similar profile for the edges connecting STG/S and PMC B (A vs. S: t = -7.09, p_FDR_ < .001; S vs. L: t = -2.81, p_FDR_ = .009; A vs. L: t = -7.58, p_FDR_ < .001). IFG/S was also coupled with PMC B, but similarly along all three sets of features (A vs. S: t = -3.63, p_FDR_ < .001; S vs. L: t = 1.02, p_FDR_ = 0.314; A vs. L: t = -1.84, p_FDR_ = .078). The language embeddings captured the most connectivity between default mode areas, such as TPJ and PMC C (A vs. S: t = -3.78, p_FDR_ < .001; S vs. L: t = -8.33, p_FDR_ < .001; A vs. L: t = -8.28, p_FDR_ < .001). Within- and between-region feature-specific model connectivity patterns were largely similar across the left and right hemispheres when examined separately (**Figs. S2, S3**). Notably, model connectivity appears to be higher in the right hemisphere for edges involving higher-order regions, particularly PMC B and to a lesser extent PMC A and C.

We also observed negative model connectivity values between the EAC and IFG/S, which may be due to systematic differences in the temporal latency of processing between these regions. In some cases, for example, between neighboring PMC B and PMC C in DMN, model connectivity was negligible, despite markedly stronger ISFC (**Fig. S4c**). Overall, these results suggest that edges connecting different language areas, including discontiguous and relatively distant cortical regions, are coupled along a subset of shared linguistic features. In line with a soft hierarchy, many connections appear to be driven by overlapping sets of linguistic features. For example, connectivity among STG/S regions is driven by acoustic, speech, and linguistic features (**Fig. 3b**). Furthermore, there are clear trends in which lower-level areas are coupled along acoustic features, and higher-level areas are coupled along language (and to a lesser extent, speech) features.

To visualize the cortical extent of feature-specific connectivity between cortical language regions, we created seed-based model connectivity maps of EAC, STG/S, and IFG/S. For each seed, we computed model connectivity for each feature band across all parcels and identified parcels with significant connectivity (using a one-sample t-test across subjects). This analysis revealed overlapping maps of model connectivity captured by each of the different feature bands for each seed (**Fig. S5**).

### Transition from acoustic- to language-based connectivity in the temporal cortex

We next zoomed in on a subset of regions for speech comprehension for which we have clear hypotheses about which linguistic features are shared across regions. We hypothesized that EAC would be coupled to neighboring STG regions primarily according to acoustic and speech features, while in more lateral STG/S regions, acoustic features of connectivity would recede and be overtaken by language features. We examined the feature-specific model connectivity between seven smaller regions along three pathways extending from the EAC/STG towards the STSva, STSvm, and STSp (**Fig. 4**).

**Fig. 4.**
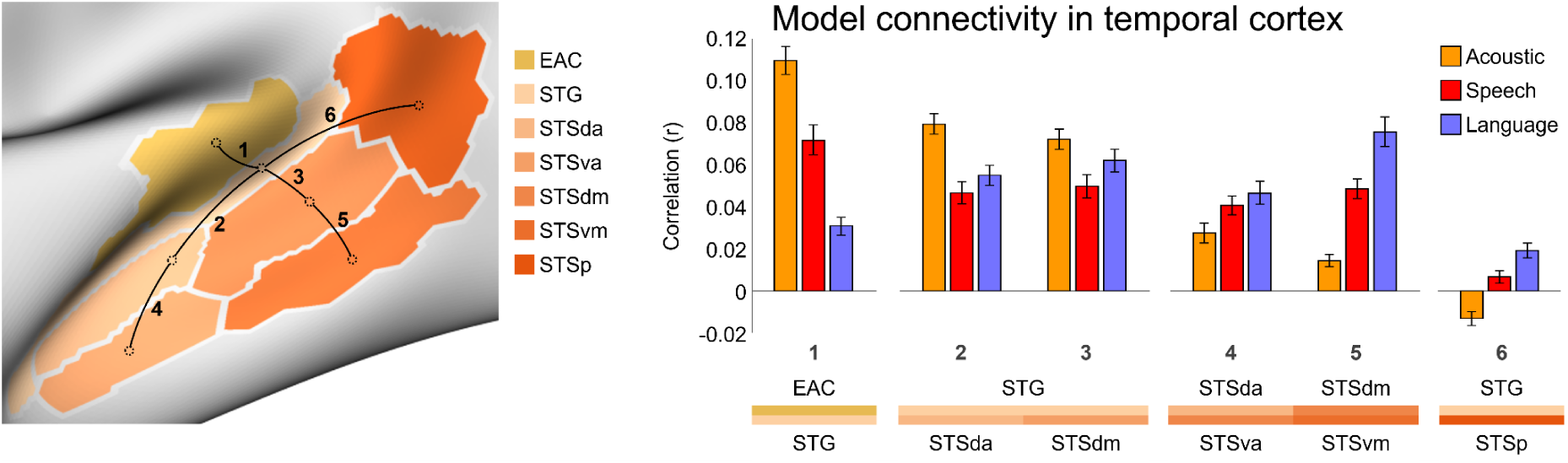
Model connectivity along superior temporal pathways. Feature-specific model connectivity was computed across parcel pairs along pathways linking early auditory cortex (EAC), superior temporal gyrus (STG), and superior temporal sulcus (STS) spanning the following regions: EAC, STG, dorsal anterior, ventral anterior, dorsal mid, ventral mid, and posterior superior temporal sulcus (STSda, STSva, STSdm, STSvm, STSp). Model connectivity showed progressive transition across feature bands as we moved further along these pathways: EAC-STG connectivity was most strongly driven by acoustic features, followed by speech features, and then language features; connectivity of STG to STSda and STSdm relatively decreased for the acoustic feature band, and all three feature bands contributed more similarly; and the pattern of connectivity from STSda to STSva and STSdm to STSvm was reversed, where language features captured the largest portion of connectivity, followed by the speech embeddings, then the acoustic embeddings. Error bars indicate bootstrap 95% confidence intervals.

We found that the model connectivity, calculated by averaging the connectivity values across all edges linking a pair of regions, progressed systematically across feature bands as we moved further along these pathways. Specifically, model connectivity between EAC and STG was most strongly driven by acoustic features, followed closely by speech features, and then language features (A vs. S: t = 12.45, p_FDR_ < .001; S vs. L: t = 11.86, p_FDR_ < .001; A vs. L: t = 20.45, p_FDR_ < .001). At the transition from STG to STSda and STSdm, acoustic connectivity relatively decreased, and all three feature bands contributed more similarly to connectivity, though some comparisons remained significant (A vs. S for STG-STSda: t = 14.02, p_FDR_ < .001; S vs. L for STG-STSda: t = -3.33, p_FDR_ = .002; A vs. L for STG-STSda: t = 7.72, p_FDR_ < .001; A vs. S for STG-STSdm: t = 9.96, p_FDR_ < .001; S vs. L for STG-STSdm: t = -4.87, p_FDR_ < .001; A vs. L for STG-STSdm: t = 3.22, p_FDR_ = .003). Finally, as we moved from STSda and STSdm to STSva and STSvm, the pattern fully reversed, such that the language features captured the largest portion of connectivity, followed by the speech embeddings, then the acoustic embeddings (A vs. S for STSda-STSva: t = -6.28, p_FDR_ < .001; S vs. L for STSda-STSva: t = -2.62, p_FDR_ = .012; A vs. L for STSda-STSva: t = -6.46, p_FDR_ < .001; A vs. S for STSdm-STSvm: t = -15.58, p_FDR_ < .001; S vs. L for STSdm-STSvm: t = -11.19, p_FDR_ < .001; A vs. L for STSdm-STSvm: t = -19.02, p_FDR_ < .001). These findings demonstrate a clear transition in the subspace of linguistic features linking EAC to increasingly lateral language areas in STG/S from predominantly acoustic to predominantly higher-level language features. Results were qualitatively similar across the left and right hemispheres when examined separately (**Fig. S6**). In all cases, however, the model-based connectivity did not fully capture stimulus-driven connectivity quantified using ISFC (**Fig. S7**).

We next devised two control analyses to assess the generalizability and specificity of our core results. First, we asked whether the observed progression from acoustic- to speech- to language-driven connectivity is specific to our choice of the Whisper model. We performed the same set of model-based connectivity analyses using acoustic and speech embeddings extracted from HuBERT and language embeddings extracted from Gemma (**Fig. S8**). We found that connectivity within and between lower-order regions were best captured by acoustic and speech features, whereas connectivity involving higher-order regions were best captured by language features—qualitatively reproducing the same soft processing hierarchy observed with Whisper. We observed some differences in performance across features when using alternative features; for example, within-EAC and EAC-STG/S connectivity was better captured by the speech embeddings rather than acoustic embeddings from HuBERT. The Whisper embeddings are theoretically superior to these ad hoc combinations of features, however, as all three levels of representation are functionally linked together in a unified model; the HuBERT and Gemma features differ in dimensionality, training diet, and learning objectives. Whisper itself provides an explicit model of how lower-level acoustic and speech features can be merged with higher-level language features in service of performing natural language tasks.

How can we ensure that our choice of acoustic, speech, and language embeddings provide a meaningful division of model features? In the second control analysis, we expressly tested these layer assignments. We performed the same set of model-based connectivity analysis after shuffling the feature assignment to the three bands before fitting the parcel-wise encoding models. While the joint model performance was unaffected (by design), the observed progression from lower-level to higher-level features was abolished by this permutation (**Fig. S9**). No differences were observed between the three permuted feature bands in capturing within- and between-region connectivity. This control analysis indicates that there is a systematic relationship between the features learned at different layers of the model and the features driving connectivity between particular areas of the language network.

### Model-based connectivity recapitulates large-scale cortical network configuration

To assess how well model connectivity captures larger-scale patterns of connectivity, we systematically examined the correlation between the feature-specific model connectivity patterns and the corresponding ISFC patterns (**Fig. S10**). Within the language network, the language embeddings better captured the ISFC pattern, followed by speech embeddings, then acoustic embeddings. For the EAC connectivity profile, acoustic model connectivity was most similar to ISFC, whereas language model connectivity was most similar to ISFC profiles for language regions (STG/S and IFG/S) and default-mode regions (TPJ and PMC). For the connectivity pattern between EAC and language regions, the acoustic and speech features best recapitulated ISFC, whereas the language features (followed by the speech features) best recapitulated the ISFC pattern between language and default-mode regions.

### Unique and shared feature-specific variance among language regions

Our analyses indicate that different regions of the language network are driven by shared subsets of stimulus features. We can also use our modeling framework to more precisely quantify what proportion of feature-specific variance in a given region is unique to that region versus shared with other regions. For a given seed region, we quantified the proportion of variance in activity captured by a given set of features that was unique to that region and the proportion that generalizes to other target regions, both relative to the joint model performance in the seed region (**Fig. 5, Table S1**). For EAC, acoustic features captured a large portion of unique variance (64%, CI: 61-66% of joint model performance), whereas a smaller portion was shared with STG/S (25%, CI: 24-27%). Acoustic and speech features corresponded to larger proportions of shared variance between EAC and STG/S than language features (acoustic: 25%, CI: 24-27%; speech: 21%, CI: 19-23%; language: 6%, CI: 5-7%). The pattern of feature-specific shared variance was reversed for the connections within STG/S regions and from STG/S to, IFG/S and PMC regions, where language features captured larger proportions of shared variance than speech and acoustic features (see **Table S1** for unique and shared proportion of joint model performance per feature set for selected seed-target pairs). For example, among STG/S regions, acoustic features captured 13% (10-15) of shared variance, whereas speech and language features each captured over twice as much shared variance (32% and 45%, respectively); similarly for connections between STG/S and IFG/S (acoustic: 8%; speech: 35%; language: 45%). Overall, this analysis revealed that higher-order regions of the language network, like STG/S and IFG/S, are characterized by larger proportions of shared variance across regions than unique, within-region variance in their activity, in contrast to EAC, particularly for higher level, more contextual features of speech and language.

**Fig. 5.**
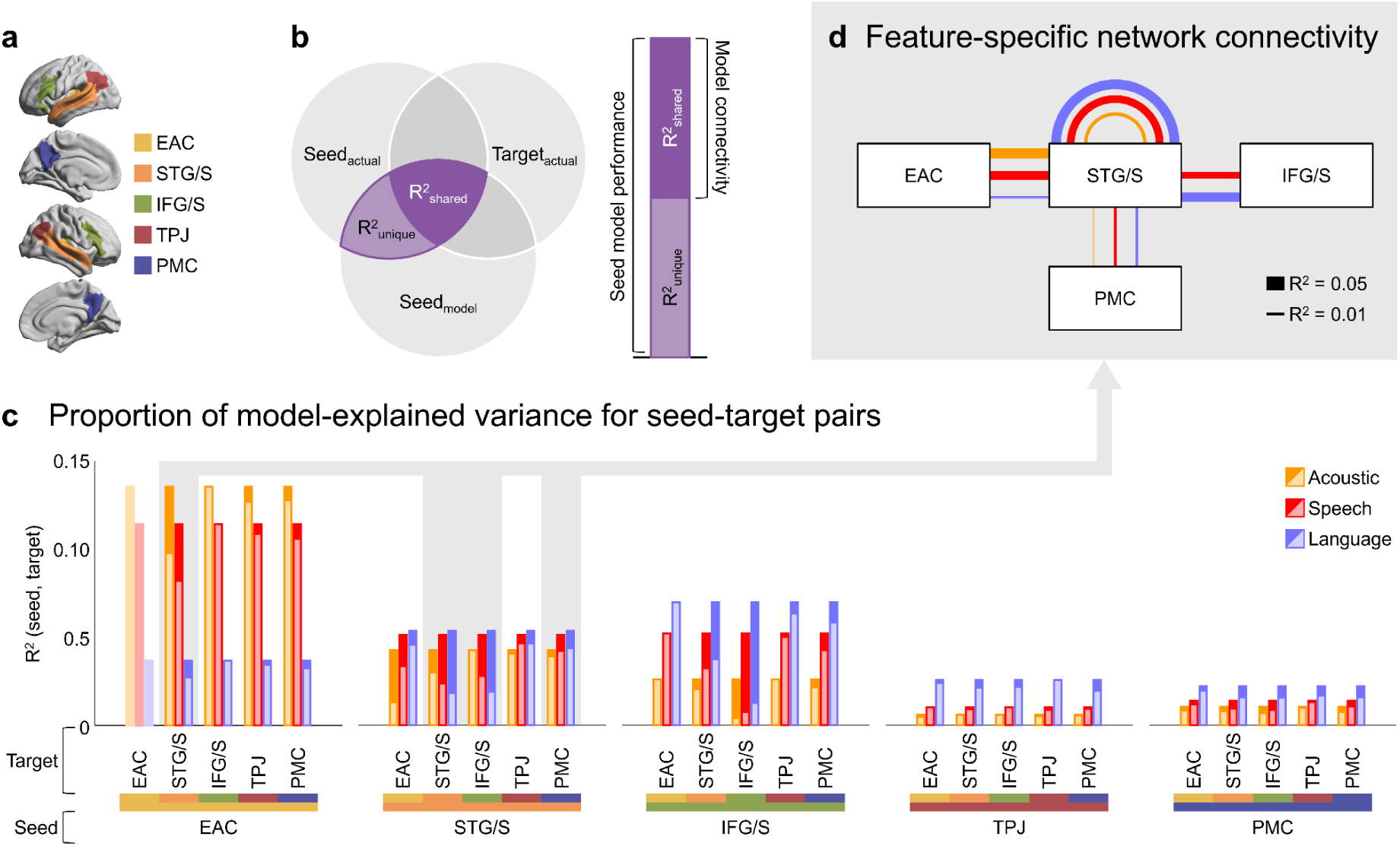
Unique and shared variance captured by encoding model among language regions. **(a)** Parcel-wise model-predicted and actual time series were averaged within 15 language regions ( **Fig. 2g**). We summarize the results within and between 5 broader regions: EAC, STG/S (consisting of STG, STSda, STSva, STSdm, STSvm, and STSp), IFG/S (IFG, IFG Orb, and IFS), TPJ (TPJ AngG, and TPJ IPL), and PMC (PMC A, PMC B, and PMC C). **(b)** The Venn diagram highlights the unique (R² unique, light purple) and shared (R² shared, dark purple) components of variance for a given pair of seed and target regions that is explained by a set of model features fit to the seed region. Note that the two components add up to the total amount of variance explained by a given feature set in the seed region (i.e., encoding performance in the seed region; corresponding to the total height of the stacked bars) and that the shared variance corresponds to model connectivity. **(c)** Unique (light color) and shared (dark color) variance for each seed-target pair is averaged across subjects and the two components are plotted as stacked bars for each feature space. Note that the EAC region contains only one parcel and therefore cannot have unique or shared variance with itself. **(d)** We highlight the amount of shared variance by feature model for a set of selected seed-target pairs along the language processing hierarchy; these pairs are highlighted in gray in the bar plot **(c)**. Each pair of regions is connected by 3 lines corresponding to the three feature models, and the line thickness corresponds to the amount of shared variance for that pair.

## Discussion

This study provides evidence that cortical language regions are coupled through distinct and overlapping subspaces of linguistic features. Using a model-based functional connectivity framework, we demonstrated that acoustic, speech, and language embeddings derived from a unified speech and language model capture distinct aspects of overall network connectivity. Acoustic features, and to a lesser extent speech features, drive connectivity between early auditory and intermediate language areas. In contrast, language features, and secondarily speech features, drive connectivity between language areas and higher-level default-mode areas. When examining more anatomically specific connectivity in superior temporal cortex, we observed a systematic transition from acoustic to language features linking neighboring temporal regions, corresponding to a processing hierarchy progressing from perceptual to more abstract features of spoken language (Kell et al., 2018). Moving from lower-level regions like EAC to higher-level language regions like STG and IFG, we also observed larger shared components of model-predicted variance across regions than unique variance to a given region, particularly for more contextualized speech and language features. Overall, these results reveal a soft processing hierarchy: regions of the language network are coupled via a mixture of acoustic, speech, and language features, but connectivity among higher-order cortical regions is driven by increasingly abstract, more contextualized features of language. This “polysemanticity” is congruent with work showing mixed selectivity for different features of natural language in individual voxels (de Heer et al., 2017) and individual neurons (Jamali et al., 2024; Khanna et al., 2024; Leonard et al., 2024), and may reflect interdependencies between acoustic and linguistic features inherent in natural speech (Wolf et al., 2024).

Over the last two decades, ISC analysis (Hasson et al., 2004; Nastase et al., 2019) has advanced the field of neuroimaging by enabling researchers to map stimulus-driven neural responses to naturalistic stimuli, like spoken language, even in the absence of an explicit model of such complex stimuli (e.g., Stephens et al., 2010; Lerner et al., 2011). In recent years, both methodological and computational advances have enabled the testing of more meaningful, neuroscientifically relevant feature representations of complex naturalistic stimuli at scale (Nastase et al., 2020a), particularly in the language domain. First, voxelwise encoding models provide a powerful framework for isolating *feature-specific* components of stimulus-driven brain activity in naturalistic paradigms (Wu et al., 2006; Naselaris et al., 2011; Wehbe et al., 2014; Huth et al., 2016, Dupré la Tour, Visconti di Oleggio Castello et al., 2025). Second, with recent advances in deep learning (e.g., Brown et al., 2020; Radford et al., 2023), we now have explicit, computational models that can accommodate the richness of real-world speech and language. Combining these advances provides a qualitative leap beyond model-free, data-driven methods like ISC, allowing researchers to test explicit models of the neural activity supporting speech, language, and communication in natural contexts (Schrimpf et al., 2021; Caucheteux et al., 2022; Goldstein et al., 2022, 2025; Tuckute et al., 2024; Zada et al., 2024).

While ISC captures stimulus-driven neural activity within a given brain region, ISFC provides a theoretical extension that enables us to quantify stimulus-driven functional connectivity *between* regions (Simony et al., 2016). Following the same logic, we evaluated our encoding models *across regions* to identify *feature-specific* functional connectivity (Toneva et al., 2022b). This model-based connectivity framework allowed us to quantify which features of speech and language drive the co-fluctuations in neural activity between regions of the language network. In the current work, we used both ISC and ISFC to quantify the upper limit of reliable, time-locked, stimulus-driven variability in the connectivity between regions (or activity within regions; i.e., a “noise ceiling”), then use encoding models to quantify how much of this stimulus-driven connectivity can be captured by specific features from a neural network model for speech and language. This approach is conceptually similar to the model connectivity method introduced by Meschke et al. (2023), which quantifies the similarity of encoding model weights across brain regions.

Our findings suggest that different areas of the language network are coupled to one another via a multidimensional space of shared linguistic features. This geometric notion of functional connectivity is loosely related to the “communication subspace” observed in visual areas by Semedo and colleagues (2019), where one population of neurons is coupled to another population through moment-to-moment co-fluctuations along a subset of dimensions in the overall activity space (Kohn et al., 2020; MacDowell et al., 2025). Geometric computations of this kind may also underlie both motor control and more abstract cognitive control (Vyas et al., 2020; Panichello & Buschman, 2021; Churchland & Shenoy, 2024); for example, preparatory motor activity can be maintained in an output-null subspace of the overall activity space to avoid prematurely triggering a motor output, then rotated into the output-potent subspace to initiate the action (Kaufman et al., 2014); similar population-level dynamics have been observed in working memory and attention tasks (Panichello & Buschman, 2021). Whisper provides an explicit, mechanistic model of how different features of speech and language can be encoded geometrically in a neural population code and merged across layers of processing in support of performing natural language tasks.

Closely related ideas have begun to emerge from efforts to understand information processing in large language models. The layers of a large language model are coupled to one another via a high-dimensional embedding space—the residual stream (Elhage et al., 2021). The attention heads (i.e., a circuit that allows the model to integrate information across words) within each layer process language by effectively reading and writing into subspaces of the residual stream. For example, an attention head in one layer can effectively communicate contextual information to an attention head in a downstream layer by modifying a subset of features in the current activity vector. In LLMs, this high-capacity residual stream is critical for encoding the rich contextual inflections of natural language; it allows the model to populate information from prior words into the activity pattern representing the current context (Desbordes et al., 2023; Muller et al., 2024). The individual “regions” of the model (i.e., the attention heads) make relatively small token-by-token adjustments to the overall activity of the shared residual stream. Different structures of language—such as phonemic, syntactic, and semantic structures—are fused into a unified embedding space, and contextualized layer by layer. In our findings, connectivity between 62% of all language parcels was dominated by contextual language features, whereas acoustic features and speech features best predicted 21% and 17% of edges, respectively. If the language network effectively integrates the contributions of different regions into a shared feature space capturing the current linguistic context, this may provide one possible computational explanation for why different regions of the language network appear to have surprisingly similar functional tuning (Fedorenko et al., 2024).

We encountered several challenges in pursuing the core questions of this work. First, we observed negative ISFC and model connectivity between certain pairs of regions, such as the connections between the EAC and DMN regions (TPJ and PMC), and the connections from TPJ to STG/S and IFG/S. For the sake of interpretational simplicity, this manuscript focuses on modeling pairs of regions with positive ISFC. We also encountered some surprising cases where the model connectivity diverged from the ISFC. For example, EAC and IFG/S were positively correlated in the ISFC matrix, but we found negative model connectivity across all three feature bands. Some of these observations could be attributed to lags in information processing between regions (Chang et al., 2022), the flexible timing incorporated into the fitting of the encoding models (Huth et al., 2016), and/or task-positive and task-negative network dynamics (Fox et al., 2005). We additionally acknowledge that the inherent spatial and temporal limitations of fMRI, as well as the potential blurring introduced by parcellation and anatomical variability across subjects, may influence the observed results and their interpretation. Finer-grained functional localization could reveal finer-grained feature selectivity within and across regions. Future work could pursue more spatial and temporally detailed analyses of interactions between regions, and how these functional interactions are supported by white-matter tracts connecting language regions (e.g., Catani et al., 2005; Saur et al., 2008; Dick et al., 2014). We resorted to relatively coarse parcel-level encoding models to reduce the computational burden and facilitate model evaluation across subjects. This choice is in tension with a body of work demonstrating that both the functional topography and functional tuning of language regions are quite variable across individual brains (Fedorenko et al., 2010; Huth et al., 2016; Mahowald & Fedorenko, 2016; Braga et al., 2020). Future work could use hyperalignment methods to obtain a finer-grained correspondence across individuals (Haxby et al., 2011, 2020; Van Uden, Nastase et al., 2018; Nastase et al., 2020b; Bhattacharjee et al., 2025).

Finally, throughout our results, we observed a sizable gap between feature-specific model connectivity and the full stimulus-driven connectivity quantified using ISFC, even when combining acoustic, speech, and language features (**Figs. S4**, **S7**). This observation raises a question: what stimulus features are driving reliable connectivity during story listening that are not yet captured by our models? This gap may ultimately shrink with larger, more densely-sampled natural language stimuli (e.g., Antonello et al., 2023; LeBel et al., 2023; Hong, Wang et al., 2024). This gap could also be due to the quality of the linguistic embeddings used to predict brain activity and connectivity. Embeddings from more advanced, and ideally more brain-like, language models may improve encoding performance and reduce the gap. The current pattern of results reveals one possible path forward. While the current selection of linguistic features captures a sizable proportion of ISFC in frontotemporal language areas, the gap is most pronounced in connections to higher-level default-mode areas (**Fig. S4c**). This suggests that current language models do not fully capture the transformation from linguistic representations to the more abstract narrative- or event-level representations thought to be encoded in the DMN (Baldassano et al., 2017; Chen et al., 2017; Yeshurun et al., 2021). As more human-like models emerge, our model-based connectivity framework provides a means to evaluate these models and reduce the gap.

## Methods

### fMRI data

This study used story-listening fMRI data from the openly available Narratives collection (Nastase et al., 2021). We used two story datasets acquired from the same 46 participants (32 females, mean age 23.33 ± 7.55). The two story stimuli are “I Knew You Were Black” (534 TRs) and “The Man Who Forgot Ray Bradbury” (558 TRs). Both stories were recorded in front of live audiences with occasional laughter, applause, and other audience reactions. “I Knew You Were Black” is an autobiographical account written and narrated by Carol Daniel which explores the intersection between her job on the radio and her identity as a Black woman. “The Man Who Forgot Ray Bradbury”, written and narrated by Neil Gaiman, is a story that explores themes of memory, forgetfulness, and language at individual and collective levels. Functional data were acquired on a 3T Siemens Magnetom Prisma with a 1.5 s TR and 2.5 mm isotropic voxels. Refer to the data descriptor for more acquisition details (Nastase et al., 2021).

### fMRI preprocessing

fMRI data were minimally preprocessed using fMRIPrep v20.0.5 (Esteban et al., 2019) including realignment, susceptibility distortion correction, spatial normalization, and resampling to the *fsaverage6* surface template (Fischl et al., 1999), as described in the data descriptor (Nastase et al., 2021). Confound regression was performed with the following nuisance variables: six head motion parameters, five principal components from both white matter and cerebrospinal fluid masks (aCompCor; Behzadi et al., 2007), cosine detrending variables, and two stimulus confounds tracking the number of words per TR, and whether a TR has words or silence. To reduce computational demands and facilitate intersubject analyses, vertex-wise time series were averaged within 1,000 parcels covering the entire cortex based on the functional atlas derived from resting-state functional connectivity (Schaefer et al., 2018).

### Regions of interest

Across both hemispheres, 280 parcels were assigned to 46 language-related regions of interest (ROIs) consisting of 23 homotopic pairs (**Fig. 2g**), defined based on four methods: functionally defined language regions (Fedorenko et al., 2010), language localizer tasks (Lipkin et al., 2022), the NeuroSynth activation map for “language” (Yarkoni et al., 2011), and intersubject correlations from 345 subjects listening to spoken stories (Nastase et al., 2021). These ROIs were selected to capture as comprehensively as possible the full cortical hierarchy for spoken language, including early auditory cortex (EAC), all core language ROIs, default-mode areas associated with event representation and narrative comprehension, as well as speech articulation areas. The procedure for defining these regions is described in detail by Zada and colleagues (2025). The 46 ROIs consisted of left and right pairs for the following regions: EAC, superior temporal gyrus (STG), dorsal anterior, ventral anterior, dorsal mid, ventral mid, and posterior superior temporal sulcus (STSda, STSva, STSdm, STSvm, STSp), inferior frontal gyrus (IFG), orbital inferior frontal gyrus (IFG orb), inferior frontal sulcus (IFS), opercular inferior frontal gyrus (IFG oper), middle frontal gyrus (MFG), superior frontal language area (SFL), supramarginal gyrus (SMG), temporoparietal junction angular gyrus and inferior parietal lobule (TPJ AngG, TPJ IPL), control C (Cont C), posterior medial cortex A, B and C (PMC A, PMC B, PMC C), parahippocampal cortex (PHC), dorsomedial prefrontal cortex (dmPFC), and sensorimotor cortex (SM). To more easily summarize our results, we also grouped 15 of these ROIs to define the following five broad, anatomically-contiguous language-related regions: EAC, superior temporal gyrus and sulcus (STG/S: STG, STSda, STSva, STSdm, STSvm, STSp), inferior frontal gyrus and sulcus (IFG/S: IFG, IFG orb, IFS, IFG oper), temporoparietal junction (TPJ: TPJ AngG, TPJ IPL), and posterior medial cortex (PMC: PMC A, PMC B, PMC C).

### Intersubject correlation and connectivity

Intersubject correlation (ISC) was computed by correlating parcel time series in each subject with the average time series across all other subjects for the corresponding parcel (i.e., leave-one-out ISC; Nastase et al., 2019). Intersubject functional connectivity (ISFC) was computed for each subject and story separately as the pairwise correlations of the subject’s parcel time series and the average parcel time series of all other subjects across all pairs of parcels. ISFC matrices were symmetrized by averaging the upper and lower off-diagonal triangles of each ISFC matrix. ISFC matrices were then averaged across the two stories. The diagonal of the ISFC matrix corresponds to the within-parcel ISC values.

Following the logic of ISC, ISFC captures stimulus-driven connectivity, because the stimulus is the only source of variance that is time-locked across subjects. Whereas traditional within-subject functional connectivity (WSFC) can be driven by intrinsic fluctuations with idiosyncratic, subject-specific timing, ISFC isolates the stimulus-driven component of connectivity (Simony et al., 2016; Simony & Chang, 2020). That said, data-driven methods like ISC, ISFC, and WSFC do not tell us what stimulus features are driving activity and/or connectivity; for example, ISC in early auditory areas may be driven by low-level acoustic features, whereas ISC in lateral temporal language areas may be driven by higher-level linguistic features. To more precisely quantify what is driving the connectivity between regions, we need to test explicit models of different stimulus features.

### Stimulus feature extraction

For the two spoken story stimuli, we extracted three types of word-level embeddings from Whisper (“openai/whisper-medium.en” from the HuggingFace library), a multimodal, transformer-based speech-to-text large language model (Radford et al., 2023). Whisper is a deep neural network using a full transformer architecture composed of separate encoder and decoder stacks. The encoder stack takes as input the speech waveform in a spectrogram format. The decoder stack takes as input text tokens corresponding to the words (or sub-words) in the audio transcript. For every word in the story, we extracted (up to) a 30-second audio segment preceding the current up until after the current word is articulated. At the same time, we extracted the words uttered in the 30-second segment from the transcript—again, ending in the current word. Then, we extracted the spectrogram from the audio, and split words into tokens. We fed this input to both the encoder and decoder in a full forward pass through the model. From the network’s activations we collected three embeddings for each word: 1) an “acoustic” embedding from the activations just prior to the first transformer layer of the encoder stack; 2) a “speech” embedding from the activations after the last layer of the encoder stack; and 3) a contextual “language” embedding from the activations of the 20th layer of the decoder stack (out of 24 layers). We use the term “acoustic” to denote that these activations are closest to the audio input; we use the term “speech” embeddings because it is the final representation of the audio—and the one that is referenced by each layer of the decoder; and we use the term “language” for the contextual word embeddings from the decoder stack because these are most similar to the embeddings extracted from typical text-based large language models (Goldstein et al., 2025). All three types of embeddings are 1,024-dimensional vectors. Timing information from the transcripts (i.e., word onsets and offsets) were used to average word-level embeddings within each corresponding fMRI TR for use in the encoding models.

To ensure the pattern of results we observed is not specific to Whisper, we reproduced our analyses using a combination of the three sets of embeddings from different models: low-level acoustic embeddings (layer 0) and higher-lever speech embeddings (layer 24) from the HuBERT speech model (“facebook/hubert-large-ll60k”; Hsu et al., 2021) and language embeddings (layer 22) from the text-based Gemma model (“google/gemma-2-9b”; Riviere et al., 2024).

To assess whether the acoustic, speech, and language embeddings extracted from Whisper capture distinct levels of representation, we permuted the assignment of features across the three types of embeddings: for a given permutation, we assigned a random third of the features to each of the (permuted) “acoustic”, “speech” and “language” feature bands. We performed 10 randomizations of this kind and averaged across permutations. This permutation effectively tests whether the division of stimulus features to acoustic, speech, and language embeddings is meaningful for predicting feature-specific connectivity between regions.

### Intersubject encoding models

We used encoding models to quantify to what extent different linguistic features are encoded in the activity of a given brain region. The two stories were used alternately for training and testing encoding models. Banded ridge regression was used to estimate parcel-wise encoding models in the train story using all three sets of embeddings (i.e., feature spaces) jointly in three separate “bands” to allow these features to fairly compete for variance in the brain activity. Each feature band was assigned its own regularization parameter based on random search across 20 log-spaced parameters in the range [1, 10^19^], using five-fold nested cross-validation within each training story. All encoding models were trained at the level of parcel time series. Encoding models of this kind quantify the average feature tuning of neural populations within each parcel. Model-predicted BOLD activity was generated for the test story based on the joint model weights, as well as for the weights at each of the three feature bands separately. Encoding models were trained within each subject, but were evaluated by correlating the model-based predictions with the actual activity at a given parcel averaged across the remaining subjects, to more closely match the formulation of the ISC analysis. In this way, all encoding models were forced to generalize both across stories and across subjects.

### Intersubject model-based connectivity

We then evaluated the encoding models fit within each parcel (from the preceding section) *across* parcels, following the logic of ISFC: the subject’s model-predicted time series was correlated with the average actual time series of all other subjects across all pairs of parcels. We refer to this analysis as intersubject model-based functional connectivity (see Toneva et al., 2022b, and Meschke et al., 2023, for related ideas). We computed a separate model-based connectivity matrix for each feature-band, as well as for the joint model. Model-based connectivity matrices were symmetrized in the same way as ISFC. When summarizing our results, we always average model-based connectivity values (correlations) among pairs of parcels, instead of averaging predicted or actual time series across parcels. ISFC conceptually serves as a noise ceiling for stimulus-driven connectivity that is reliable across subjects. In implementation, however, given that encoding models are more flexible in accounting for hemodynamic lags, model-based connectivity may numerically diverge from ISFC values.

In summarizing model connectivity results, we focused on eight language-related regions comprising 207 parcels that have been shown to exhibit strong ISC during story listening: EAC, STG/S, IFG/S, TPJ AngG, TPJ IPL, PMC A, PMC B, and PMC C. Within-region (model) connectivity was summarized for a given region by averaging the correlation values between all pairs of parcels within that region separately for each subject and feature band. Between-region (model) connectivity between a given pair of regions was summarized by averaging the correlation values between all pairs of parcels connecting the two regions. For visualization and interpretational simplicity, we opted not to evaluate model performance across regions (i.e., model connectivity) for pairs of regions with negative ISFC values.

### Unique and shared model-predicted variance

We next aimed to quantify the proportions of model-predicted variance that are unique to a given (seed) region versus shared with other (target) regions. Encoding models were first fit and evaluated within parcels. For a given seed region, encoding performance quantifies the total variance in a seed region explained by the model (or a subset of features), whereas model-based connectivity quantifies the variance shared between the model-based predictions for the seed region and a target region. That said, some of the variance explained by a model in a seed region may be either unique to that seed or shared with a target parcel. To quantify the proportions of model-explained variance unique to a given seed region versus shared with a target region, we apply a semi-partial correlation method to model-predicted and actual time series for the seed and target regions (computed by averaging across all parcel-level time series within the seed and target regions separately). Specifically, to quantify the explained variance unique to a given seed region (R^2^ unique) in a given subject, we first regressed the target region’s time series (averaged across other subjects) out of the seed time series, then computed the R^2^ between that subject’s model-predicted time series and the residual time series for the seed region (also averaged across other subjects). To quantify the total variance shared between the seed and target captured by the seed model, we compute the variance explained (R^2^ total) by a subject’s seed model time series in the seed actual time series (averaged across other subjects). The shared variance (R^2^ shared) is then quantified as the difference between R^2^ total and R^2^ unique. We then normalized these unique and shared R^2^ values into proportions of the R^2^ total for the joint model, so that proportions for each feature space are compared to the same reference. These percentages are not intended to sum to one, but rather indicate proportions of the joint model performance. Note that proportions may not be symmetric between seed and target (e.g., EAC-STG/S vs. STG/S-EAC); one region may encode a subset of features that are encoded in another region, or may have a different signal-to-noise ratio.

### Statistical testing

We assessed the statistical significance of whole-brain ISC and encoding model performance at each parcel using a one-sample *t*-test across subjects (**Fig. 1**). The false discovery rate (FDR) was controlled at *p* < .05 across 1,000 parcels. To determine whether model-based connectivity values differed significantly from one another, we performed paired *t*-tests (*df* = 45) between the model performance values for different feature bands. FDR was controlled at *p* < .05 among the comparisons under consideration. For visualization, we generated error bars by bootstrapping subject-level values (i.e., resampling subjects with replacement) 1,000 times for each mean value.

## Data, Materials, and Software Availability

The fMRI data used here are openly available as part of the Narratives collection (Nastase et al., 2021): https://doi.org/10.18112/openneuro.ds002345.v1.1.4; https://datasets.datalad.org/?dir=/labs/hasson/narratives; https://fcon_1000.projects.nitrc.org/indi/retro/Narratives.html. The code used to perform the core analyses of this study is available at https://github.com/zaidzada/narrative-enc.

## Acknowledgments

We are grateful for the use of data from the freely-available Narratives collection. We would like to acknowledge funding sources: UBC Friedman Award for Scholars in Health and BC Children’s Hospital Research Institute Doctoral Studentship (AS); National Institutes of Health CRCNS grant R01DC022534 (ZZ, UH, SAN).

## Author contributions

Conceptualization: AS, ZZ, UH, SAN

curation: ZZ, SAN

Formal analysis: AS, ZZ

Funding acquisition: AS, TV, UH

Investigation: AS, ZZ

Methodology: AS, ZZ, SAN

Project administration: SAN

Software: AS, ZZ

Supervision: TV, UH, SAN

Visualization: AS

Writing – original draft: AS, ZZ, SAN

Writing – review & editing: AS, ZZ, TV, UH, SAN

## Supplementary Information

**Fig. S1.**
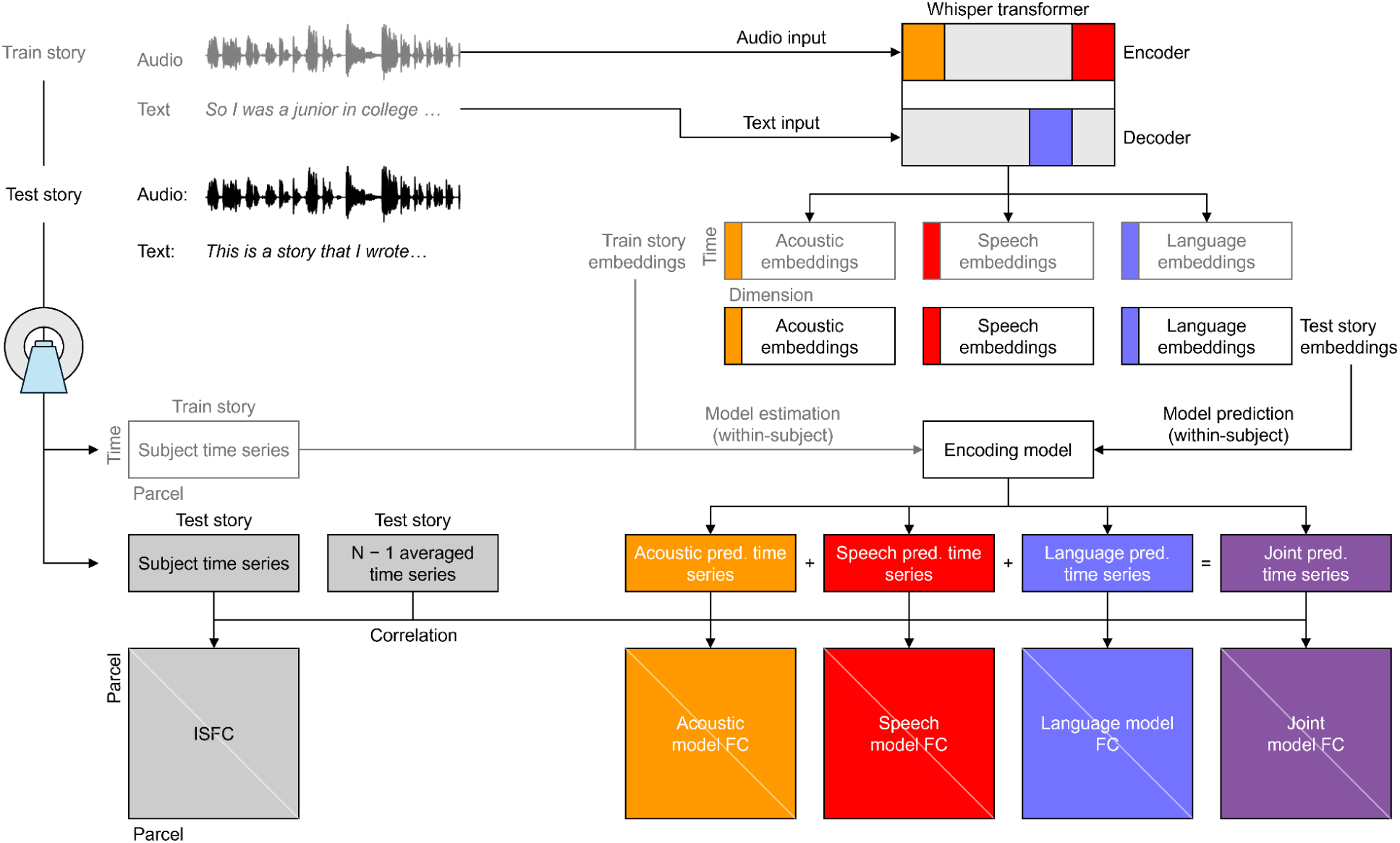
Schematic workflow and encoding model evaluation. fMRI data were collected from 46 subjects while they listened to two spoken narratives: each story served as the train story and test story. The transcript and audio spectrograms of the train and test stories were fed into a large language model (Whisper) to extract distinct word-level embedding representations of the story as follows: acoustic embeddings were extracted from the input to the first transformer layer of the encoder stack; speech embeddings were extracted from the last encoder stack; and language embeddings were extracted layer 20 of the decoder stack. The train story embeddings were then used in combination with train story parcel-resolution fMRI time series data to estimate parcel-wise encoding models using banded ridge regression. Model-based predictions of parcel time series were generated based on the weights associated with each of the three feature bands, as well as the joint model weights. In parallel, intersubject function connectivity (ISFC) was computed as the pairwise correlation of a given subject’s time series and the group-averaged time series of all other subjects for all pairs of brain parcels. Model-based connectivity was also computed following the logic of ISFC by correlating the subject’s predicted time series and the group-average actual time series of all other subjects for all pairs of parcels.

**Fig. S2.**
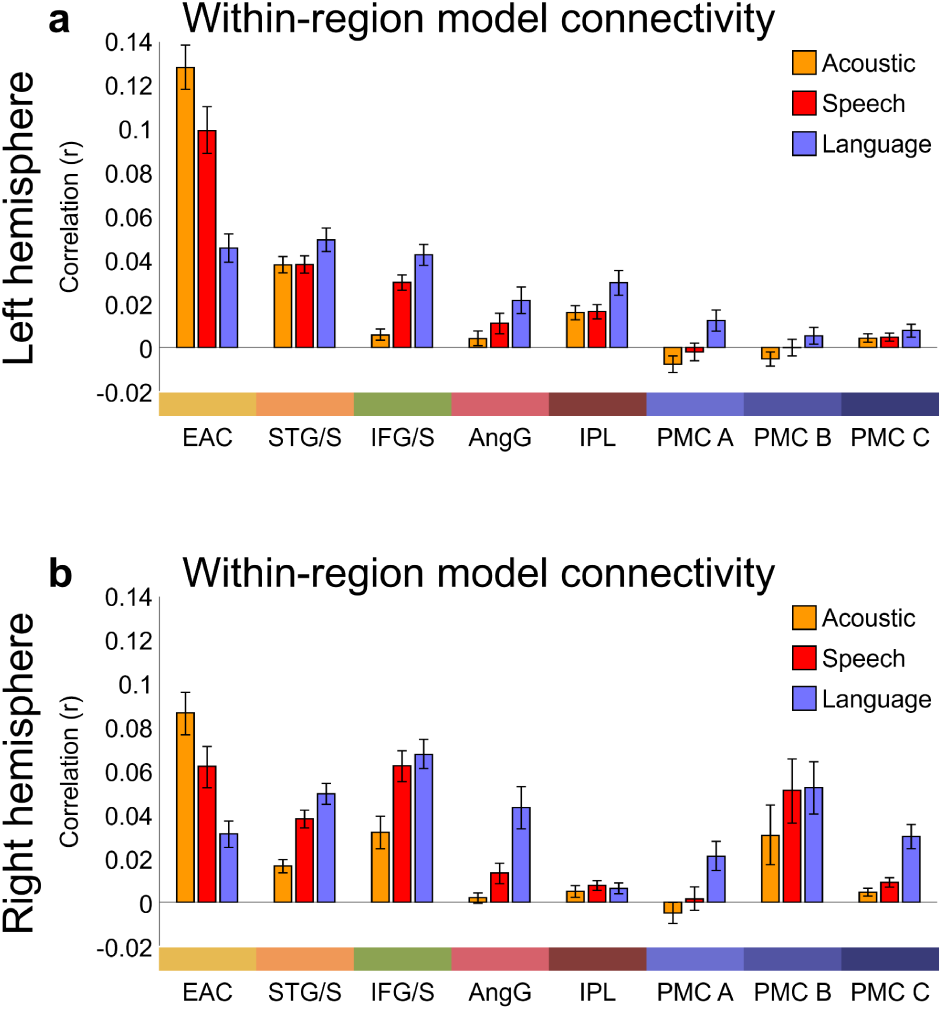
Within-region feature-specific model connectivity plotted separately for the left and right hemispheres, related to Fig. 3. Feature-specific model connectivity values corresponding to **Fig. 3b** are shown separately for the left **(a)** and right **(b)** hemispheres. Connectivity values were averaged across parcel pairs within each language region for each hemisphere independently. Error bars indicate bootstrap 95% confidence intervals.

**Fig. S3.**
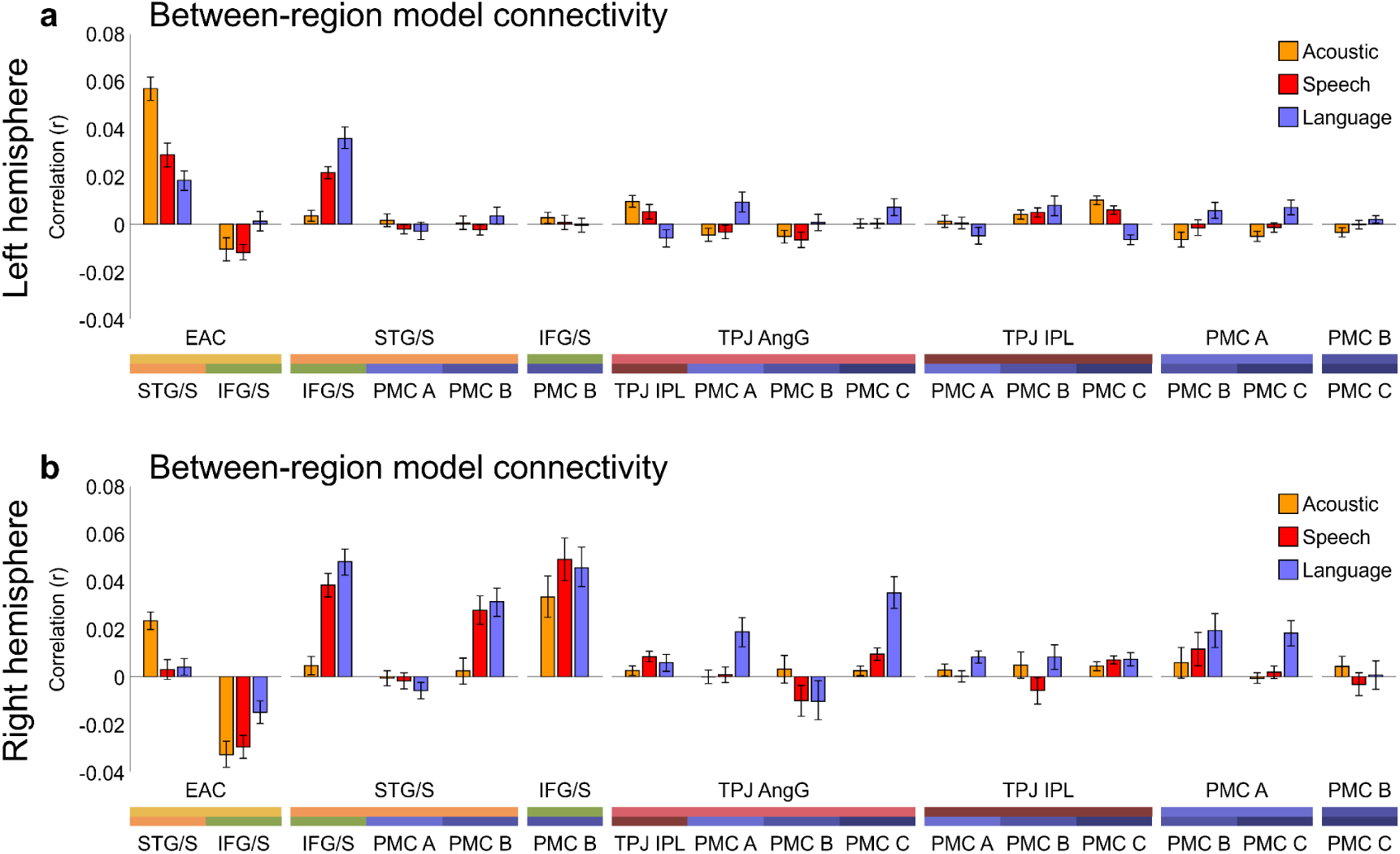
Between-region feature-specific model connectivity plotted separately for the left and right hemispheres, related to Fig. 3c. Feature-specific model connectivity values corresponding to **Fig. 3c** are shown separately for the left **(a)** and right **(b)** hemispheres. Connectivity values were averaged across parcel pairs connecting different language regions within each hemisphere. Between-region model connectivity results are shown only for region pairs with positive ISFC values. Error bars indicate bootstrap 95% confidence intervals.

**Fig. S4.**
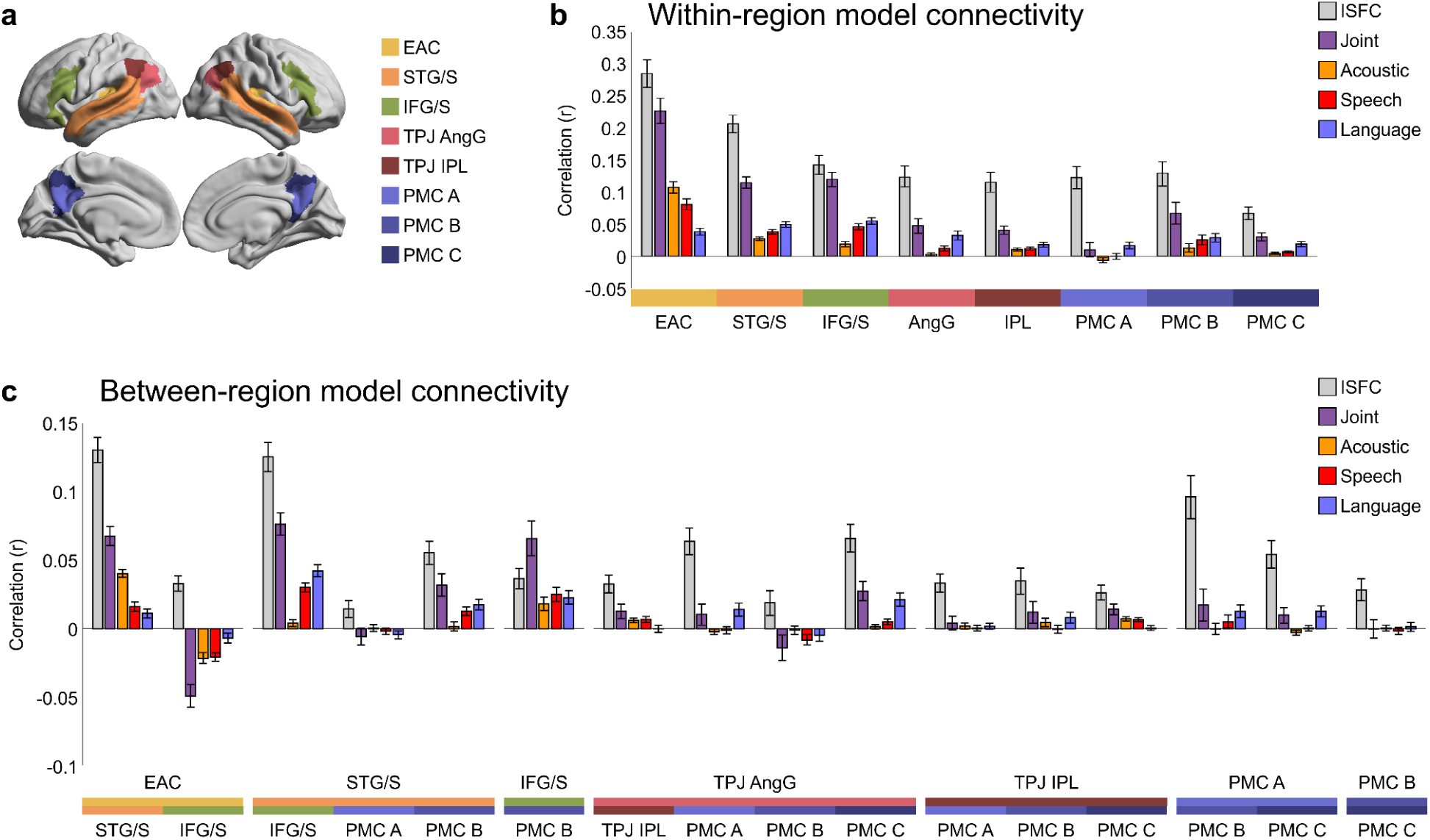
Model-based connectivity within and between language regions in reference to intersubject functional connectivity, related to Fig. 3. **(a)** Focusing on the same eight language-related regions (early auditory cortex [EAC], superior temporal gyrus and sulcus, [STG/S], inferior frontal gyrus and sulcus [IFG/S], temporoparietal junction angular gyrus and inferior parietal lobule [TPJ AngG and TPJ IPL], and posterior medial cortex A, B and C [PMC A, PMC B, and PMC C]) from **Fig. 3**, we reproduced the feature-specific model connectivity results with the addition of the joint model connectivity and intersubject functional connectivity (ISFC) values for both the within **(b)** and between **(c)** language region. Similar to **Fig. 3**, between-region model connectivity results are shown only for region pairs with positive ISFC values. Error bars indicate bootstrap 95% confidence intervals.

**Fig. S5.**
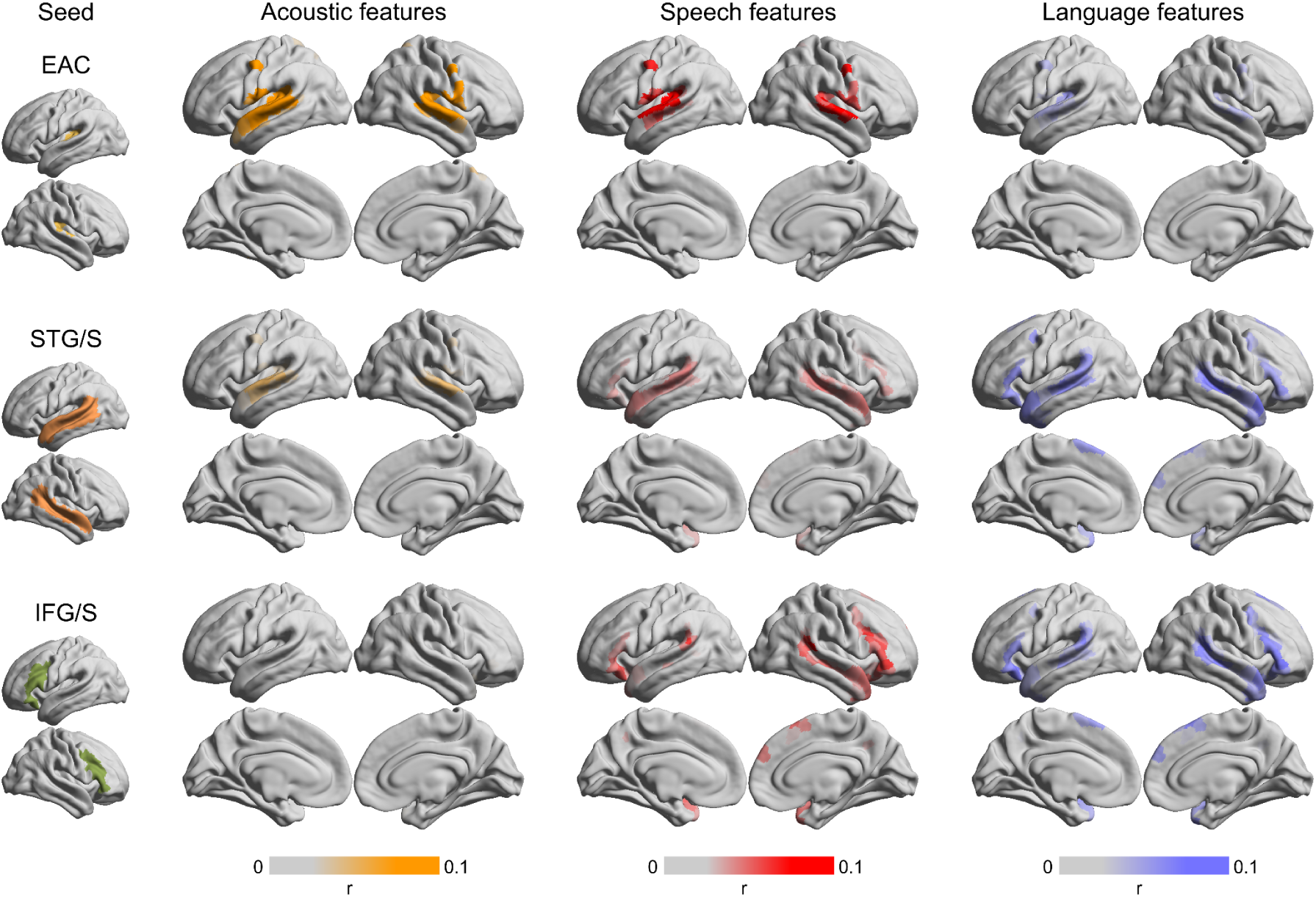
Seed-based feature-specific connectivity maps. Seed-based model connectivity maps were computed separately for the three feature bands in three seed regions: early auditory cortex (EAC), superior temporal gyrus and sulcus (STG/S), and inferior frontal gyrus and sulcus (IFG/S). For each feature band and seed, we first computed the parcel-pair connectivity between parcels within the seed and all parcels with significant encoding based on the joint model (i.e., the same regions as in **Fig. 1e**). Then, we averaged the resulting connectivity values across the seed parcels. Models trained in EAC yielded significant feature-specific model connectivity maps constrained to perisylvian areas, with the acoustic embeddings driving the marginally wider-spread connectivity. For models trained in STG, connectivity based on acoustic embeddings was tightly localized to the middle STG, whereas speech embeddings yielded connectivity extending along the STG/S, and language embeddings expanded connectivity in the STG/S and IFG/S. For models trained in IFG/S, only weak encoding was found for acoustic embeddings, whereas the speech and language embeddings yielded increasingly widespread connectivity in frontotemporal language areas. This suggests that language areas are coupled through multiple, overlapping sets of linguistic features. However, higher-level linguistic features (speech and language embeddings) are increasingly dominant in linking higher-order regions.

**Fig. S6.**
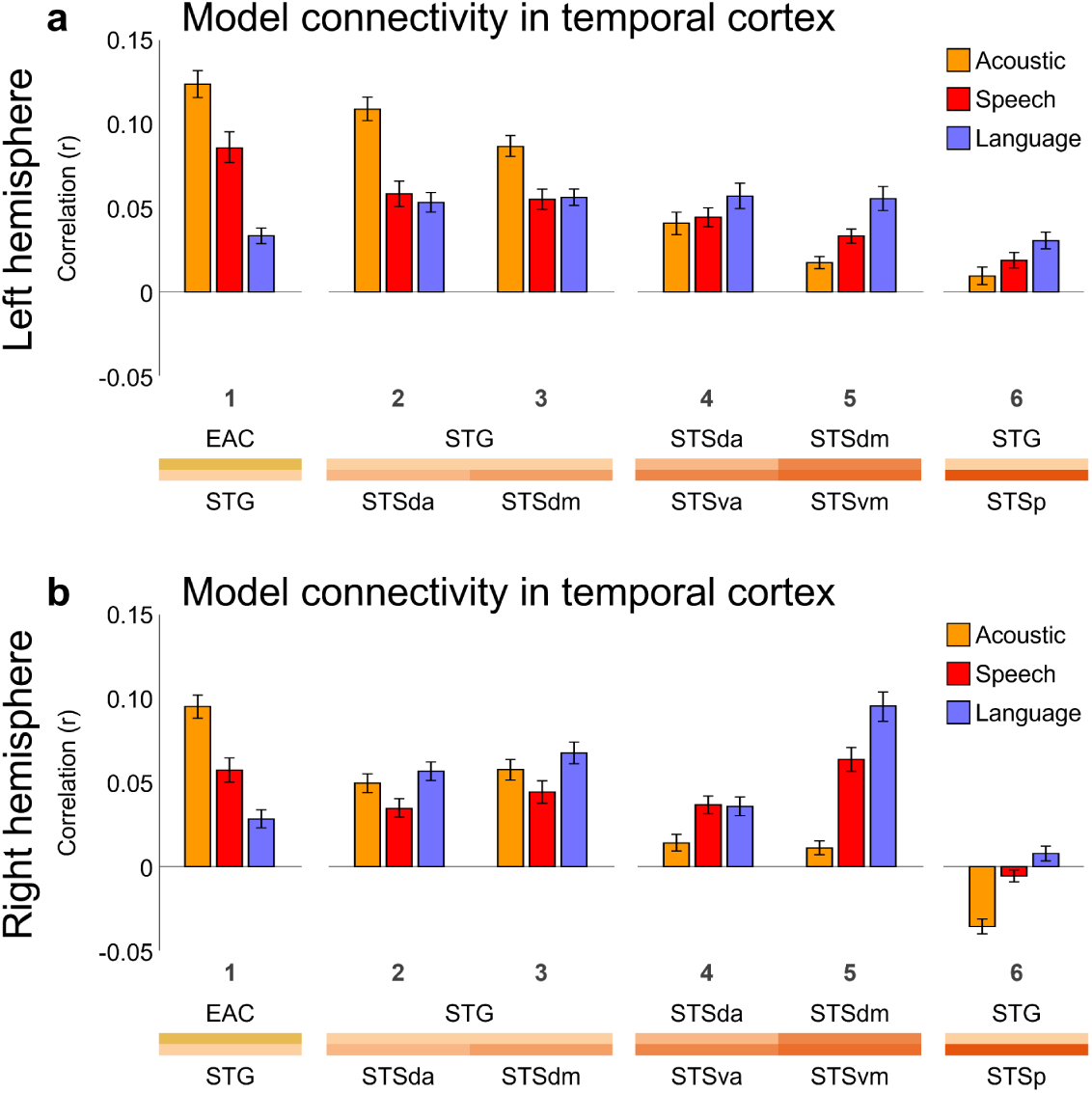
Model connectivity along superior temporal pathways plotted separately for the left and right hemispheres, related to Fig. 4. Feature-specific model connectivity corresponding to **Fig. 4** is shown separately for the left **(a)** and right **(b)** hemispheres. Model connectivity was computed across parcel pairs along pathways linking early auditory cortex (EAC), superior temporal gyrus (STG), and superior temporal sulcus (STS), spanning EAC, STG, dorsal anterior, ventral anterior, dorsal mid, ventral mid, and posterior STS (STSda, STSva, STSdm, STSvm, STSp). The qualitative progression in connectivity captured by each feature along these pathways is preserved across hemispheres. Error bars indicate bootstrap 95% confidence intervals.

**Fig. S7.**
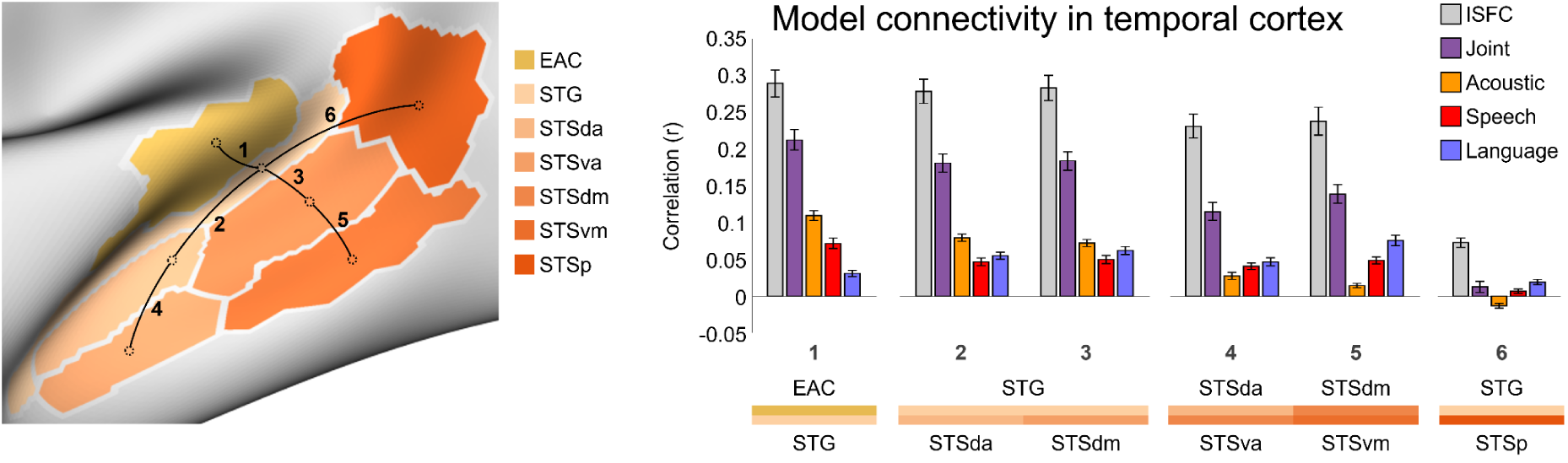
Model connectivity along superior temporal pathways in reference to intersubject functional connectivity, related to Fig. 4. We reproduced model connectivity results across parcel pairs along pathways linking early auditory cortex (EAC), superior temporal gyrus (STG), and superior temporal sulcus (STS) with the addition of joint model connectivity and intersubject functional connectivity (ISFC) values for reference. Error bars indicate bootstrap 95% confidence intervals.

**Fig. S8.**
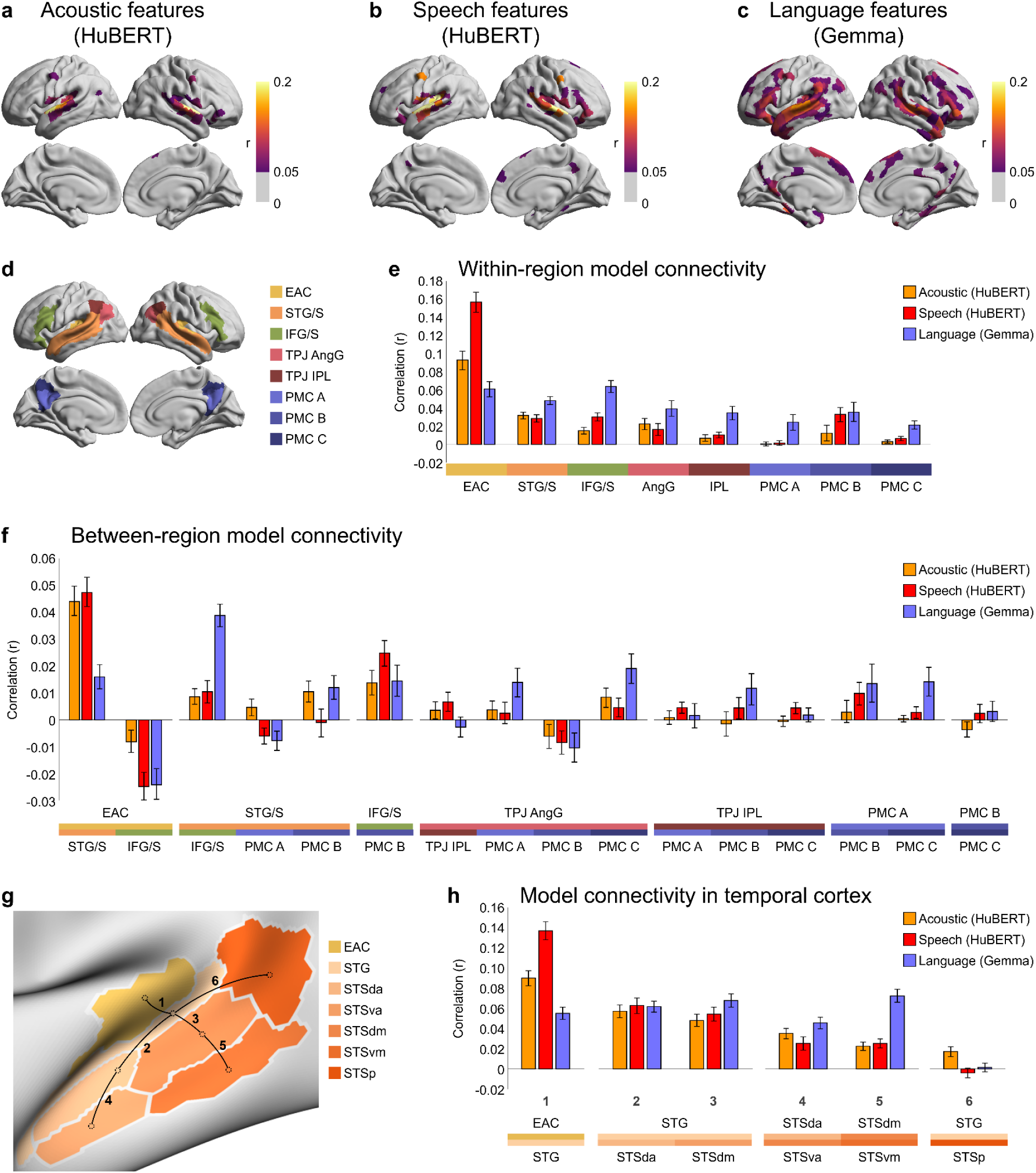
Feature-specific encoding performance and model connectivity using embeddings extracted from alternative models (HuBERT and Gemma). **(a–c)** Results corresponding to **Fig. 1g**, showing feature-specific model performance using acoustic and speech embeddings extracted from HuBERT and linguistic embeddings extracted from Gemma. Encoding models and evaluation procedures are identical to those used in **Fig. 1**. **(d–f)** results corresponding to Fig. 3, showing feature-specific model-based connectivity within language regions **(e)** and between language regions **(f)**, computed using HuBERT acoustic and speech embeddings and Gemma linguistic embeddings. Connectivity values were averaged across parcel pairs within and between language regions, and between-region results are shown only for region pairs with positive ISFC values. **(g–h)** Results corresponding to **Fig. 4**, showing model connectivity along superior temporal pathways linking early auditory cortex (EAC), superior temporal gyrus (STG), and superior temporal sulcus (STS). Connectivity was computed across parcel pairs spanning EAC, STG, dorsal anterior, ventral anterior, dorsal mid, ventral mid, and posterior STS (STSda, STSva, STSdm, STSvm, STSp) using HuBERT and Gemma embeddings. The qualitative progression of feature-specific model connectivity lower-order to higher-order regions is preserved. Error bars indicate bootstrap 95% confidence intervals.

**Fig. S9.**
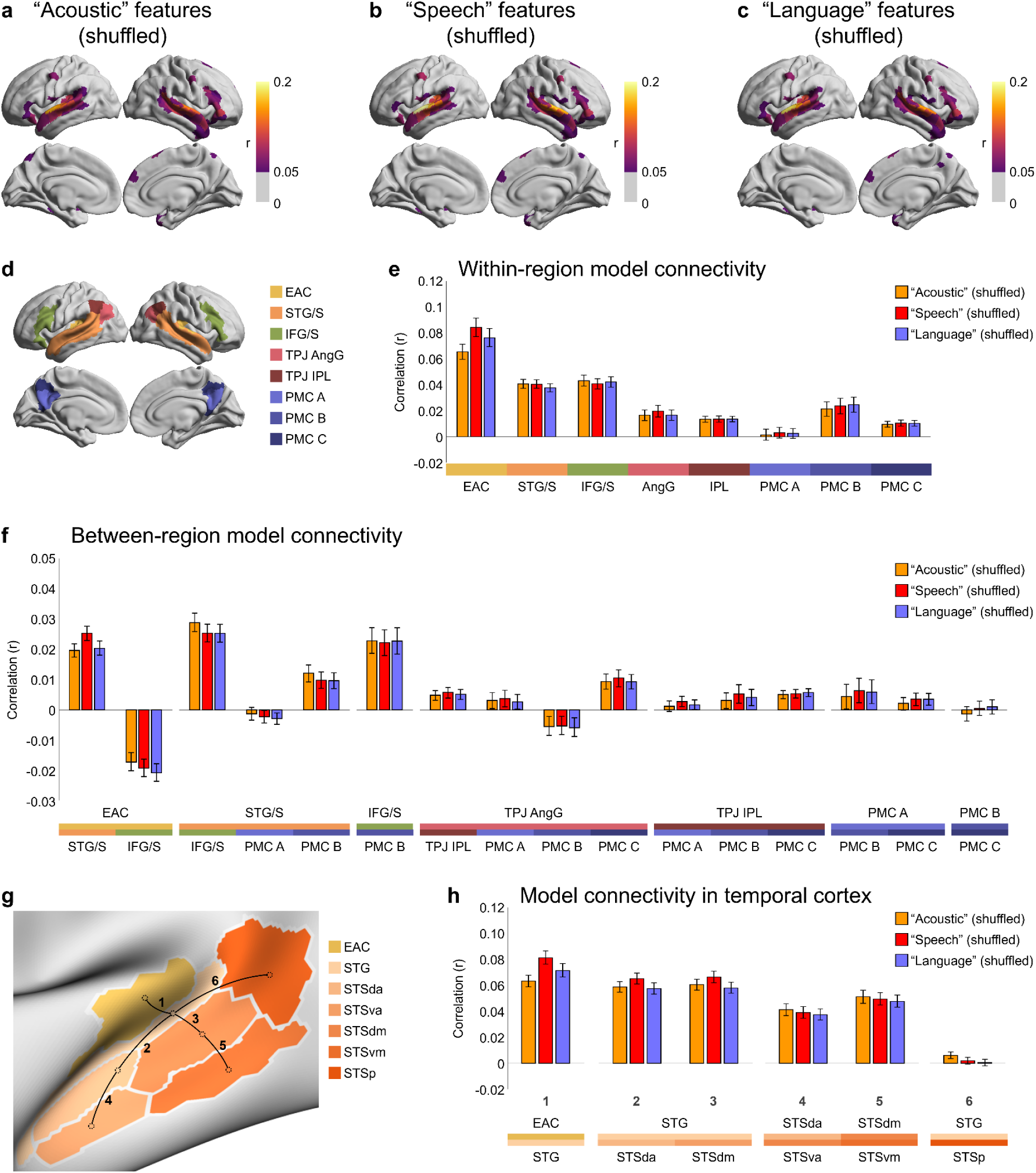
Feature-specific encoding performance and model connectivity: control analysis with randomly shuffled feature-band assignments. **(a–c)** Results corresponding to **Fig. 1g**, shown for encoding models using Whisper embeddings with the assignment of features to the acoustic, speech, and language bands randomly shuffled. Model construction, fitting, and evaluation are otherwise identical to the main analysis. **(d–f)** Results corresponding to **Fig. 3**, showing feature-specific model-based connectivity within and between language regions computed using Whisper embeddings with randomly shuffled feature-band assignments. Connectivity values were averaged across parcel pairs within and between language regions, and between-region results are shown only for region pairs with positive ISFC values. **(g–h)** Results corresponding to **Fig. 4**, showing model connectivity along superior temporal pathways computed using Whisper embeddings with shuffled feature-band assignments. In all cases under this control manipulation, differences between feature bands in capturing within-and between-region connectivity are largely abolished, indicating that the feature-specific connectivity effects observed in the main analyses depend on the structured assignment of features to bands. Error bars indicate bootstrap 95% confidence intervals.

**Fig. S10.**
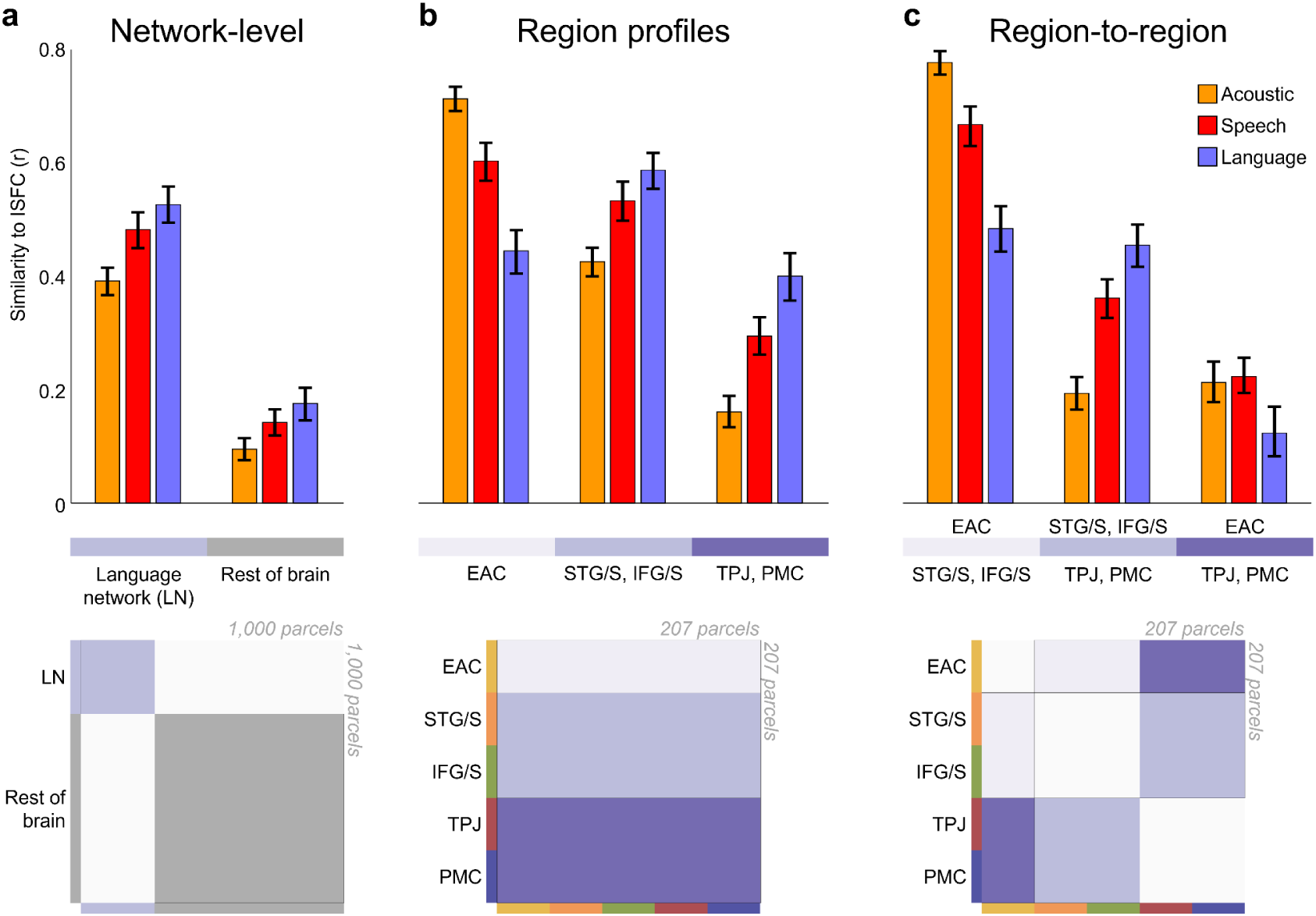
The feature-specific model connectivity recapitulates the intersubject functional connectivity network configuration. The similarity between model connectivity and intersubject functional connectivity (ISFC) was quantified as the correlation between vectorized model connectivity and ISFC (sub)matrices. The highlighted (purple) parts in the schematic parcel-by-parcel connectivity matrices at the bottom indicate the edges contributing to each group of bars. **(a)** Similarity between patterns of feature-specific model connectivity and ISFC was computed within the whole language network and, as a control, within the rest of the cortex. Within the whole language network, the language embeddings better captured the ISFC pattern, followed by speech embeddings, then acoustic embeddings (A vs. S: t = -7.09, p_FDR_ < .001; S vs. L: t = -4.11, p_FDR_ < .001; A vs. L: t = -7.81, p_FDR_ < .001). Across the rest of the cortex, the similarity between the feature-specific model connectivity patterns and ISFC was much lower for all features (A in language network vs. rest of brain: t = 25.98, p_FDR_ < .001; S in language network vs. rest of brain: t = 26.44, p_FDR_ < .001; L in language network vs. rest of brain: t = 29.90, p_FDR_ < .001). **(b)** Similarity between model connectivity and ISFC was computed for three broad groups of regions associated with spoken narrative comprehension across their respective connectivity profiles: early auditory cortex (EAC), language (i.e., superior temporal gyrus and sulcus [STG/S] and inferior frontal gyrus and sulcus [IFG/S], and default-mode (i.e., TPJ and posterior medial cortex [PMC]) regions. For EAC connectivity profile, acoustic model connectivity was most similar to ISFC (A vs. S for EAC regions: t = 8.00, p_FDR_ < .001; S vs. L for EAC regions: t = 8.46, p_FDR_ < .001; A vs. L for EAC regions: t = 14.89, p_FDR_ < .001), whereas language model connectivity was most similar to ISFC profiles of language (STG/S and IFG/S) and default-mode (TPJ and PMC) regions (A vs. S for language regions: t = -7.57, p_FDR_ < .001; S vs. L for language regions: t = -4.68, p_FDR_ < .001; A vs. L for language regions: t = -9.09, p_FDR_ < .001; A vs. S for default-mode regions: t = -9.51, p_FDR_ < .001; S vs. L for default-mode regions: t = -6.81, p_FDR_ < .001; A vs. L for default-mode regions: t = -11.07, p_FDR_ < .001). **(c)** Finally, we examined the similarity of model connectivity and ISFC between groups of low-, mid, and high-level regions associated with spoken narrative comprehension. For the connectivity pattern between EAC and downstream frontotemporal language areas, the acoustic and speech features best recapitulated ISFC (A vs. S: t = 7.59, p_FDR_ < .001; S vs. L: t = 10.39, p_FDR_ < .001; A vs. L: t = 15.42, p_FDR_ < .001), whereas the language features (followed by the speech features) best recapitulated the ISFC pattern between language and default-mode areas (A vs. S: t = -12.17, p_FDR_ < .001; S vs. L: t = -5.88, p_FDR_ < .001; A vs. L: t = -12.49, p_FDR_ < .001). The similarity between model connectivity and ISFC was fairly similar for acoustic and language features, with speech features slightly (but significantly) outperforming the other features for the connections between EAC and default-mode areas (A vs. S: t = -0.56, p_FDR_ = .058; S vs. L: t = 5.29, p_FDR_ < .001; A vs. L: t = 3.92, p_FDR_ < .001). Error bars indicate bootstrap 95% confidence intervals.

**Table S1.** Unique and shared model-predicted variance within and between language regions along the language processing hierarchy, related to Fig. 5. This table summarizes the proportions of feature-specific variance that is unique to a seed region (R^2^ unique) or shared with another target region (R^2^ shared) and normalized by the seed model variance for selected seed-target pairs. The 95% confidence interval (CI) for each percentage (based on 1,000 bootstraps resampling subjects) is provided between parentheses.

| Seed →<br>target | % $R^2$ unique (CI) | | | % $R^2$ shared (CI) | | |
| --- | --- | --- | --- | --- | --- | --- |
|  | Acoustic | Speech | Language | Acoustic | Speech | Language |
| EAC → STG | 64% (61-66) | 54% (50-58) | 19% (16-21) | 25% (24-27) | 21% (19-23) | 6% (5-7) |
| STG → STG | 34% (32-37) | 30% (28-32) | 28% (26-30) | 13% (10-15) | 32% (30-35) | 45% (42-48) |
| STG → IFG | 39% (36-42) | 28% (25-30) | 28% (25-31) | 8% (6-11) | 35% (32-38) | 45% (42-49) |
| STG → PMC | 41% (38-43) | 49% (47-52) | 61% (58-63) | 6% (5-7) | 13% (12-14) | 12% (11-14) |

